# Evolution of crab eye structures and the utility of ommatidia morphology in resolving phylogeny

**DOI:** 10.1101/786087

**Authors:** Javier Luque, W. Ted Allison, Heather D. Bracken-Grissom, Kelsey M. Jenkins, A. Richard Palmer, Megan L. Porter, Joanna M. Wolfe

**Affiliations:** Department of Geology and Geophysics, Yale University, New Haven, CT 06520-8109, USA; Department of Biological Sciences, University of Alberta, Edmonton, Alberta T6G 2E9, Canada; Smithsonian Tropical Research Institute, Balboa–Ancón 0843–03092, Panamá, Panamá; Department of Biological Sciences, Florida International University-Biscayne Bay Campus, North Miami, FL 33181, USA; Department of Biology, University of Hawaii at Manoa, Honolulu, HI 96822, USA; Museum of Comparative Zoology and Department of Organismal and Evolutionary Biology, Harvard University, Cambridge, MA 02138, USA

**Keywords:** Apposition, Brachyura, Cretaceous, Cenozoic, exceptional preservation, fossil, Superposition

## Abstract

Image-forming compound eyes are such a valuable adaptation that similar visual systems have evolved independently across crustaceans. But if different compound eye types have evolved independently multiple times, how useful are eye structures and ommatidia morphology for resolving phylogenetic relationships? Crabs are ideal study organisms to explore these questions because they have a good fossil record extending back into the Jurassic, they possess a great variety of optical designs, and details of eye form can be compared between extant and fossil groups. True crabs, or Brachyura, have been traditionally divided into two groups based on the position of the sexual openings in males and females: the so-called ‘Podotremata’ (females bearing their sexual openings on the legs), and the Eubrachyura, or ‘higher’ true crabs (females bearing their sexual openings on the thorax). Although Eubrachyura appears to be monophyletic, the monophyly of podotreme crabs remains controversial and therefore requires exploration of new character systems. The earliest podotremous lineages share the plesiomorphic condition of ‘mirror’ reflecting superposition eyes with most shrimp, lobsters, and anomurans (false crabs and allies). The optical mechanisms of fossil and extant podotreme groups more closely related to Eubrachyura, however, are still poorly investigated. To better judge the phylogenetic utility of compound eye form, we investigated the distribution of eye types in fossil and extant podotreme crabs. Our findings suggest the plesiomorphic ‘mirror’ eyes—seen in most decapod crustaceans including the earliest true crabs—has been lost in several ‘higher’ podotremes and in eubrachyurans. We conclude that the secondary retention of larval apposition eyes has existed in eubrachyurans and some podotremes since at least the Early Cretaceous, and that the distribution of eye types among true crabs supports a paraphyletic podotreme grade, as suggested by recent molecular and morphological phylogenetic studies. We also review photoreceptor structure and visual pigment evolution, currently known in crabs exclusively from eubrachyuran representatives. These topics are critical for future expansion of research on podotremes to deeply investigate the homology of eye types across crabs.

## INTRODUCTION

True crabs, or Brachyura, are a speciose and economically important group of crustaceans first known from the Early Jurassic, more than 170 Mya (Luque et al., 2019a; Wolfe et al., 2019, and references therein). Their remarkable modern and past diversity of form and adaptations is not restricted to their carapace or limbs (Fig. 1), but also evident in the wide range of compound eye types and underlying visual systems across modern crab taxa (Cronin and Porter, 2008) (Fig. 2). Brachyura are widely recognized as a monophyletic clade (e.g., Rice, 1981; Jamieson et al., 1995; Ahyong et al., 2007; Ng et al., 2008; Scholtz and McLay, 2009; Tsang et al., 2014; Luque et al., 2019a; Wolfe et al., 2019). Yet, their internal relationships are still debated, especially concerning the so-called ‘podotreme’ crabs (i.e., those crabs where both males and females have coxal sexual openings, Fig. 3A), and how they relate to Eubrachyura or ‘higher’ crabs, with sexual openings on the coxa of males and the thorax in females (i.e., Heterotremata, Fig. 3B), or thoracic in both males and females (i.e., Thoracotremata, Fig. 3C).

**Figure 1.**
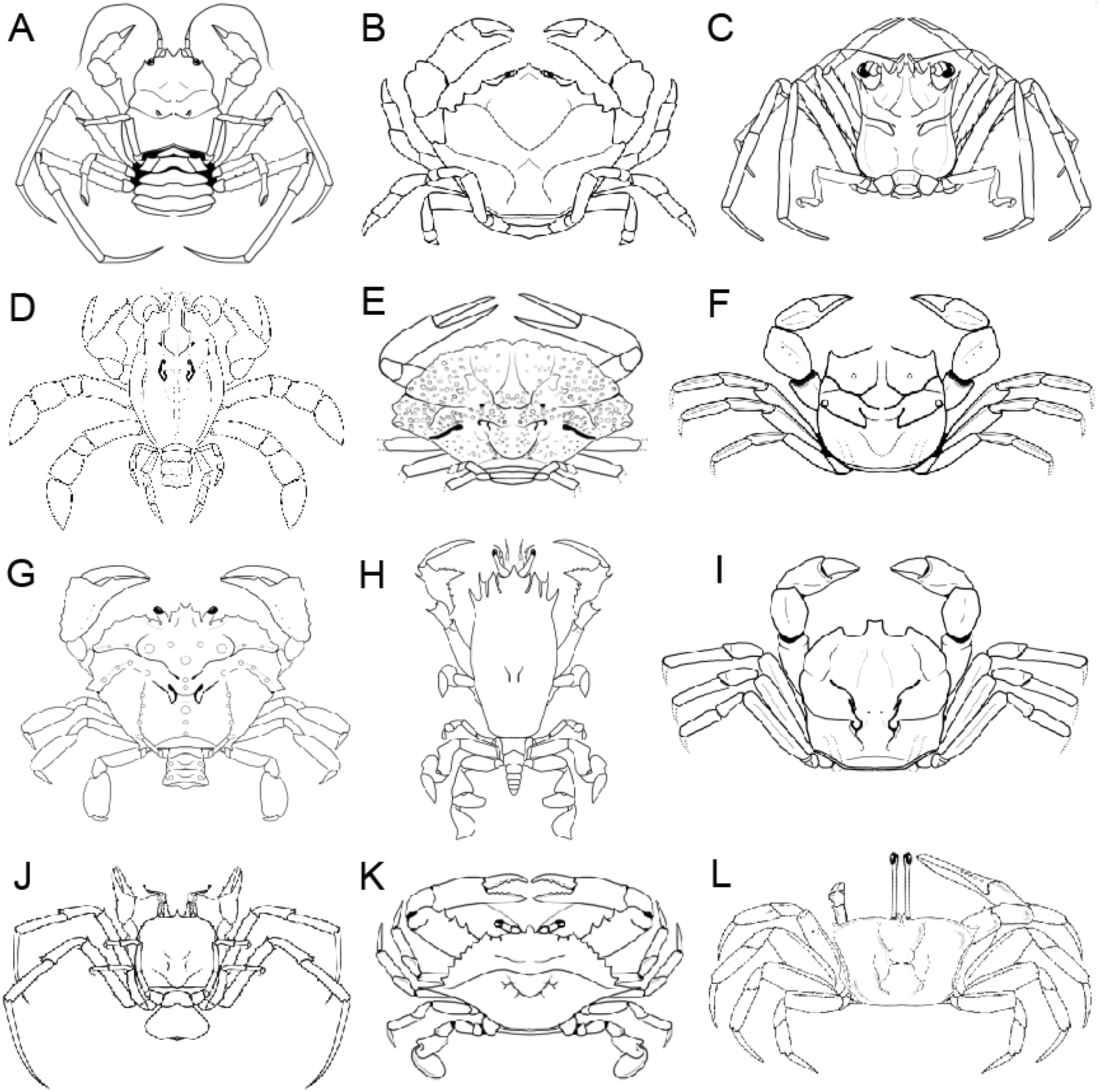
Diversity of form across the main extant and fossil groups of true crabs. A, Homolodromioidea. B, Dromioidea. C, Homoloidea. D, †Callichimaeroidea. E, †Etyoidea. F, †Torynommoidea. G. Raninoida: †Necrocarcinoidea. H, Raninoidea. I, †Dakoticancroidea. J. Cyclodorippoidea. K, Eubrachyura: Heterotremata. L, Eubrachyura: Thoracotremata. Dagger (†) indicates extinct groups (modified from Luque et al., 2019a). Illustrations and figure by J. Luque. Line drawings not to scale.

**Figure 2.**
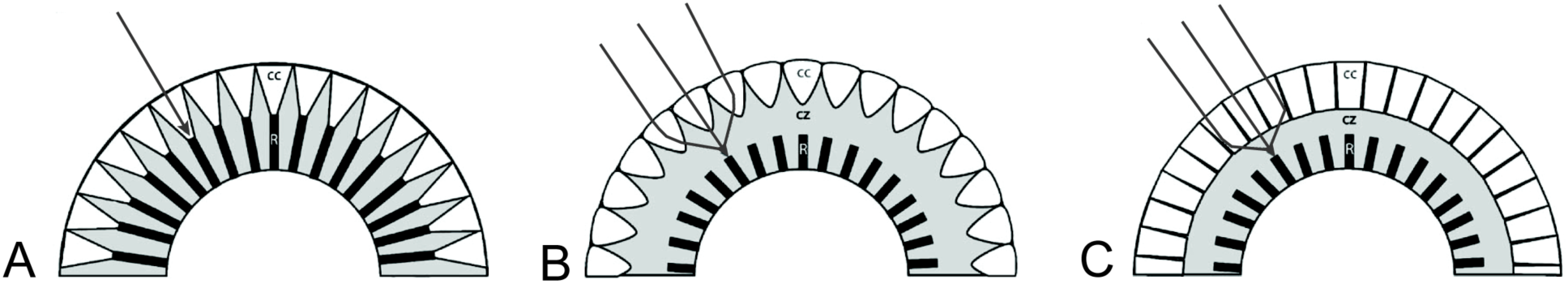
Types of compound eyes in brachyuran crabs. A: <u>Apposition eye</u>-works well in relatively bright light. A single light beam is focused on the retina of a single ommatidium (thin line). B–C: superposition eyes are better suited for vision in dim light. Recognized by the presence of a “clear zone” (grey area) between the outer lenses (white) and the retina (black). B: <u>Refracting superposition eye</u>-the crystalline cones contain a refractive index gradient that bends incoming light to focus it on the retina (thin lines); multiple light rays may fall on a single ommatidial retina. C: <u>Reflecting superposition eye</u>-light rays are focused by reflections off the sides of the cones, which are square instead of round in cross section, and are typical of ‘mirror’ optics; here too, multiple beams of light may fall on a single ommatidial retina. Abbreviations: cc=crystalline cone; cz=clear zone; r=rhabdom. Drawings modified from Cronin and Porter (2008). Parabolic superposition not illustrated.

**Figure 3.**
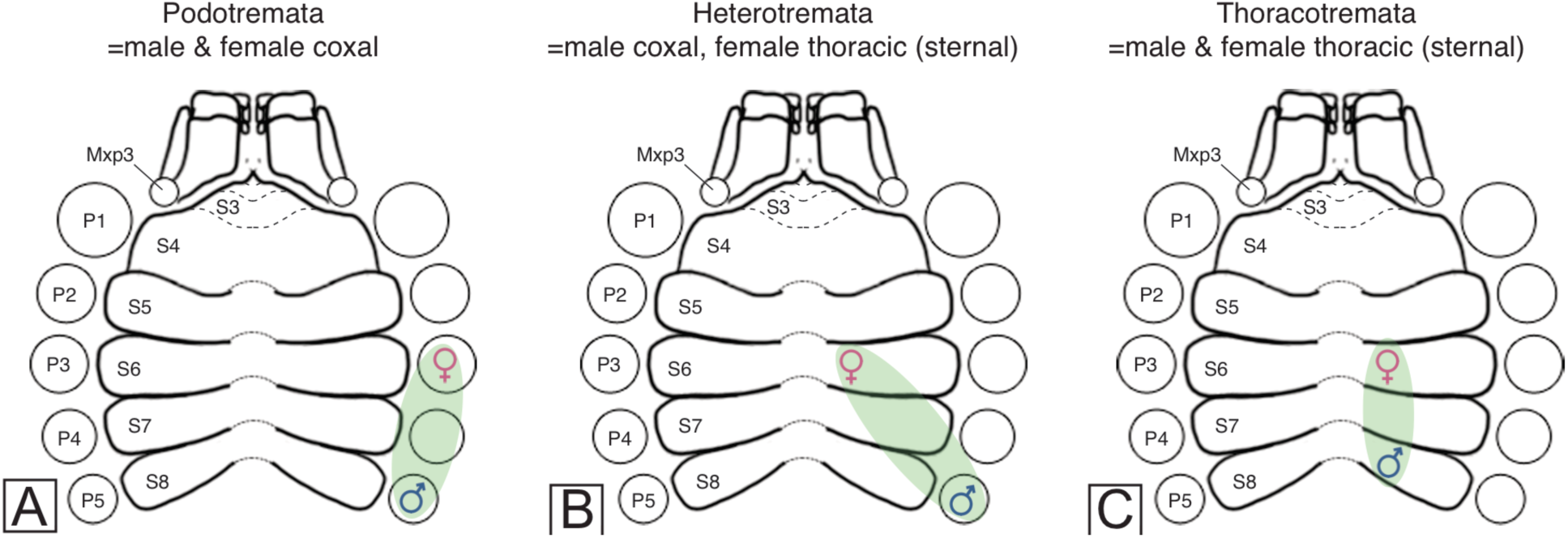
Schematic diagram showing the position of sexual openings in brachyuran crabs. A, podotreme condition; B, heterotreme condition; C, thoracotreme condition. For all decapods, including Anomura and Brachyura, the plesiomorphic condition is males and females with coxal sexual openings, or podotreme. The innovation of sternal sexual openings in female crabs is presumed to have occurred once in the most recent common ancestor for heterotreme and thoracotreme crabs (=Eubrachyura). Illustrations and figure by J. Luque.

The lack of agreement about how podotremes relate to each other and to eubrachyurans has profound effects on our understanding of the evolution of true crabs (Luque et al., 2019a), and therefore any potential phylogenetic significance that a given eye type, and therefore its facet shape and packing, may have (Fig. 4). Although closely related groups would be expected to share similar visual systems and facet shapes/packing, no work has investigated the distribution of eye types across crabs in a phylogenetic context to date, neither whether the morphology of their ommatidia is useful to infer underlying eye types, and especially, what does the fossil record tell about the distribution of facet shape and packing across crabs through time.

**Figure 4.**
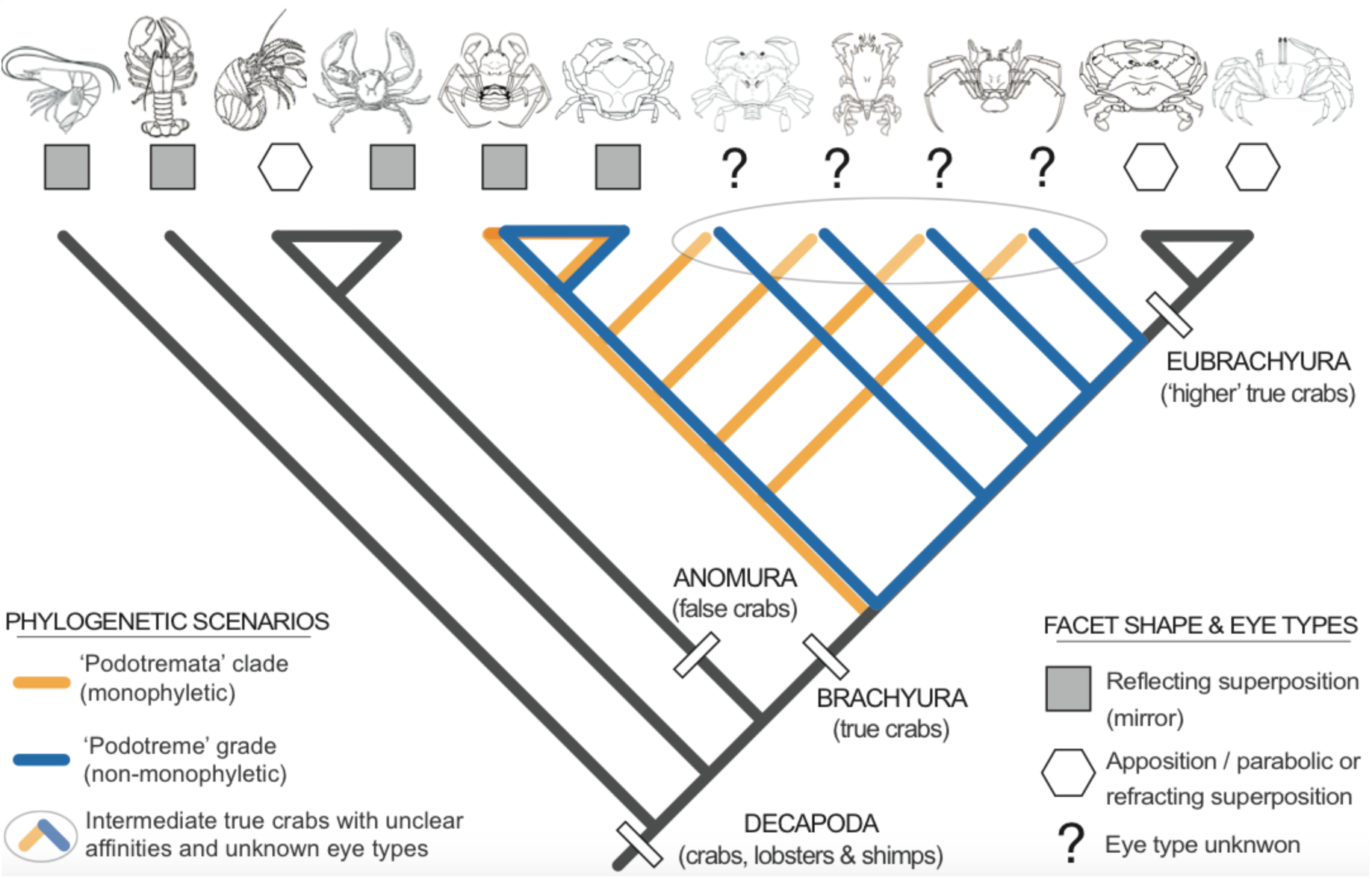
Schematic phylogenetic scenarios for the evolution of true crabs (Brachyura), and the distribution of their facet shape, packing, and visual systems. The main competing hypotheses suggest that either ‘lower’ true crabs, or podotremes, form a monophyletic clade Podotremata (orange lines), whereas podotreme crabs may represent a paraphyletic grade of increasing complexity (blue lines) with some intermediate groups closer to eubrachyurans than to other podotremes. ‘Lower’ brachyurans share reflecting superposition ‘mirror’ eyes (grey squares), while the ‘higher’ brachyurans lack mirror eyes altogether and have either apposition, parabolic superposition, or refracting superposition eyes (white hexagons). In this work, we investigate the visual systems in intermediate ‘lower’ podotremes (white oval, and marked with ‘**?**’), to assess the utility of ommatidia morphology in resolving crab phylogeny. Figure by J. Luque.

### Basic eye types in crabs

Currently, there are four main types of compound eyes recognized in crustaceans: apposition, parabolic superposition, refracting superposition, and reflecting superposition, each one with its particular combination of external and internal features (Land, 1976; Nilsson, 1988; Gaten, 1998; Cronin and Porter, 2008) (Fig. 2). Apposition eyes are the simplest (Fig. 2A). In this eye type, isolated ommatidia with hexagonal facets are packed in a hexagonal lattice, and functions best in relatively bright light. They are the ancestral condition for crustaceans and present in the larval stages of all decapods (Land, 1980; Fincham, 1984; Porter and Cronin, 2009) (Fig. 5).

**Figure 5.**
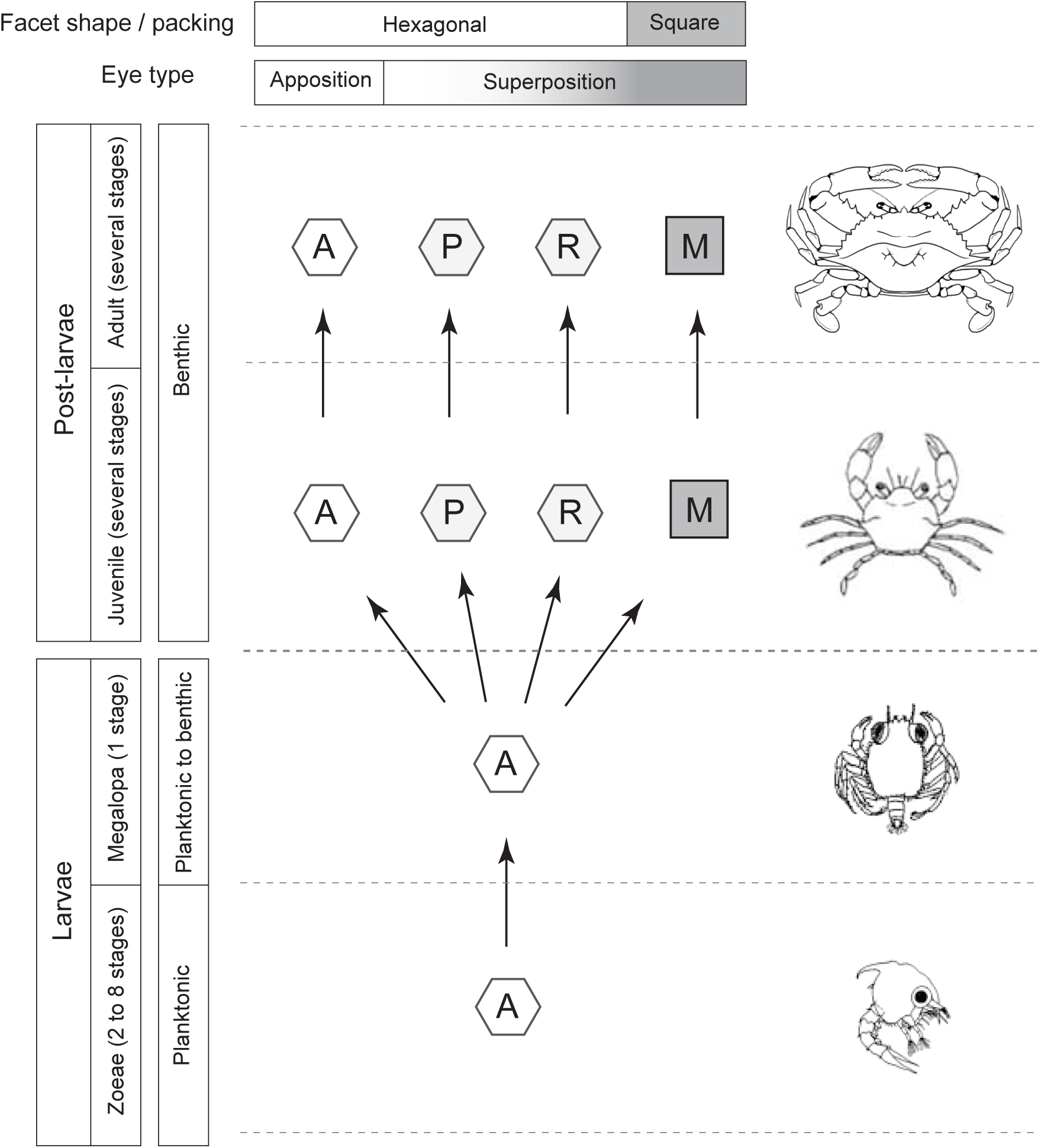
Distribution of eye types in crabs across life stages. The larval stages of brachyurans and other decapod crustaceans have apposition eyes, which is the ancestral state for malacostracans. In post-larval stages, the larval apposition eyes may either remain functional as apposition eyes, or undergo internal and external restructuration to function as superposition eyes. Externally, apposition (A), parabolic superposition (P), and refracting superposition (R) eyes share the hexagonal packing of hexagonal to round facets, while eyes of the reflecting superposition type (M) are modified to work as a mirror box, and have square facets with orthogonal packing. Figure by J. Luque.

The other three eye types are of the superposition type, which are better–suited for vision in dim light condition, and differ from apposition eyes by the presence of a “clear zone” between the outer structures of the eye and the retina (Cronin and Porter, 2008) (Fig. 2B–C). Parabolic superposition eyes predominantly have hexagonal facet shapes in hexagonal packing, while the sides of the crystalline cones (the structure under each facet of the eye surface) are shaped in the form of a parabola and the ommatidia have a light guide that focuses the collimated light onto the retina (Fincham, 1980; Nilsson, 1989). Refracting superposition eyes (Fig. 2B) also have hexagonal facets in hexagonal packing, but their crystalline cones have a refractive index gradient that bends incoming light to focus it on the retina (Nilsson et al., 1986; Nilsson, 1990). Finally, the reflecting superposition eyes (Fig. 2C) lack the refractive index gradient of the refracting superposition eye or the light guides of the parabolic superposition type, but instead focus an image by reflecting light off the sides of the crystalline cones as occurs in a four-sided mirror box, hence the common name “mirror eyes” (Vogt, 1975; Land, 1976). Unlike the facets of apposition, parabolic superposition, and refracting superposition eyes, which share the presence of hexagonal to roundish facets packed in a hexagonal lattice, reflecting superposition eyes have distinctive square facets packed in an orthogonal lattice and are also square in cross-section (Fig. 5). Among crustaceans, mirror eyes are unique to decapods, and are the only eye type with rectangular, square facets.

Interestingly, while most crustaceans groups with compound eyes show only the apposition type through larval and adult stages, crabs alone have representatives of all four eye types (Porter and Cronin, 2009) (Fig. 5). The evolutionary history of apposition and superposition eyes is still poorly understood (Nilsson, 1983; Gaten, 1998). In particular, little is known about the genetic and developmental mechanisms regulating the expression of a particular eye type in the post-larva, and the information provided by the fossil record has been sparse and fragmentary, until now.

### Eye types in larval and post-larval crabs

Most larval and adult crustaceans, including crabs, have compound eyes of the apposition type (Gaten, 1998) (Fig. 5), suggesting that this eye type is the ancestral condition for crustaceans. However, among crustaceans, reflecting superposition or ‘mirror’ eyes are unique to post-larval Decapoda (Land, 2000). They are present in most extant penaeoid and caridean shrimp, lobsters, anomurans such as Galetheoidea (squat lobsters) and some pylochelideans (symmetrical hermit crabs), and the podotreme brachyurans Dromioidea, Homolodromioidea and Homoloidea (Gaten, 1998; Porter and Cronin, 2009; Scholtz and McLay, 2009). As such, the absence of mirror eyes in several derived crab groups is intriguing. In Eubrachyura or ‘higher’ crabs, the loss of reflecting superposition optics via secondary retention of larval apposition eyes appears to have occurred in their most recent common ancestor by progenetic paedomorphosis (Gaten, 1998).

Based solely on the position of sexual openings, taxonomists have traditionally grouped true crabs into Podotremata, Heterotremata, and Thoracotremata (Guinot, 1977) (Fig. 2). This taxonomic grouping presumes that a) Heterotremata and Thoracotremata are monophyletic (together forming the section Eubrachyura), and b) Podotremata, or the ‘lower’ Brachyura, are monophyletic and form the sister group to Eubrachyura. However, the podotreme condition of coxal sexual openings is plesiomorphic and shared with all anomurans, other decapods, and even heterotreme brachyuran males, casting doubts on its utility for classifying crab taxa (Luque et al., 2019a and references therein). Moreover, most studies dealing with phylogenetic analyses have recovered a paraphyletic podotreme grade (e.g., Ahyong et al., 2007; Scholtz and McLay, 2009; Karasawa et al., 2011; Tsang et al., 2014; Luque et al., 2019a; Wolfe et al., 2019). In addition, the distribution of visual systems across brachyuran clades is poorly understood. Early branches of crown podotremes like Homoloidea, Dromiodea, and Homoloidea have ‘mirror’ eyes—which are plesiomorphic for crown Decapoda—while adult eubrachyurans have reverted to larval apposition eyes (Gaten, 1998). Therefore, the most derived group of crabs exhibits the most plesiomorphic eye morphology. Yet, almost nothing is known about the eye types present in ‘intermediate’ podotreme groups, either fossil or extant (Fig. 4), which has motivated the present study.

As Gaten (1998) suggested, if the stratigraphic ranges of the fossil and extant decapod crustacean groups can be combined with information about their eye types, then some phylogenetic patterns may appear. Here, we present a comprehensive review on the visual systems of true crabs, and integrate novel data on the external eye features of fossil and extant brachyurans. As very little is known about the eyes in fossil and living ‘intermediate’ crabs, we explore the extents and limitations of using eye form as an additional tool to understand crab evolution, while providing an overview of the ecology and development of crab visual systems, and their phylogenetic implications.

## MATERIALS AND METHODS

### Institutional abbreviations

**AMNH**: American Museum of Natural History, New York, USA.
**IGM p**: Colecciones Paleontológicas Museo José Royo y Gómez, Servicio Geológico Colombiano, Bogotá D.C., Colombia.
**MNHN**: Muséum national d’Histoire naturelle, Paris, France.
**MUN-STRI**: Mapuka Museum of Universidad del Norte, Barranquilla, Colombia.
**NPL**: Non-vertebrate Paleontology Lab, Jackson School Museum of Earth History, University of Texas, USA.
**QMW**: Queensland Museum, Brisbane, Australia.
**USNM**: United States National Museum of Natural History, Smithsonian Institution, Washington, D.C., USA.
**YPM**: Invertebrate Zoology Collections, Yale Peabody Museum, Yale University, New Haven, Connecticut, USA.

### Materials

#### Extant taxa

Nineteen extant species across all podotreme superfamilies were studied from the invertebrate zoology collections of the USNM, MNHN, and QMW. Specimens were preserved in 70% EtOH, and one eye from selected adult specimens was removed for microscope imaging and preserved in 70%EtOH. Illustrated taxa included *Dicranodromia felderi* Martin, 1990 (Homolodromioidea: Homolodromiidae) (Fig. 6A–C); *Dromia personata* (Linnaeus, 1758) (Fig. 6E–F) and *Hypoconcha* sp. (Fig. 6G–I) (Dromioidea: Dromiidae), *Dynomene filhol* Bouvier, 1894 (Dromioidea: Dynomenidae) (Fig. J–L); *Homola minima* Guinot and Richer de Forges, 1995 (Fig. 7A–C), and *Latreillopsis bispinosa* Henderson, 1888 (Fig. 7D–G) (Homoloidea: Homolidae), *Eplumula phalangium* (De Haan, 1839) (Homoloidea: Latreillidae) (Fig. 7H–J); *Lysirude nitidus* (A. Milne-Edwards, 1880) (=*Lyreidus bairdii*) (Fig. 8A–C) and *Lysirude griffini* Goeke, 1985 (Fig. 8D–F) (Raninoidea: Lyreididae), *Cyrtorhina granulosa* Monod, 1956 (Fig. 8G–I), *Symethis* sp. (Fig. J–L), *Cosmonotus grayi* White, 1848 (Fig. 9A–C), *Notopus dorsipes* (Linnaeus, 1758) (Fig. 9D–F), *Ranilia muricata* H Milne Edwards, 1837 (Fig. 9G–I), *Ranina ranina* (Linnaeus, 1758) (Fig. J–L), *Notopoides latus* Henderson, 1888 (Fig. 10A–C), *Notosceles viaderi* Ward, 1942 (Fig. 10D–F), *Raninoides benedicti* Rathbun, 1935b (Raninoidea: Raninidae) (Fig. 10G–I); and *Clythrocerus nitidus* (A. Milne-Edwards, 1880) (Cyclodorippoidea: Cyclodorippidae) (Fig. 10J–L). A complete list of the extant material studied and associated information is provided in Table 1.

**Figure 6.**
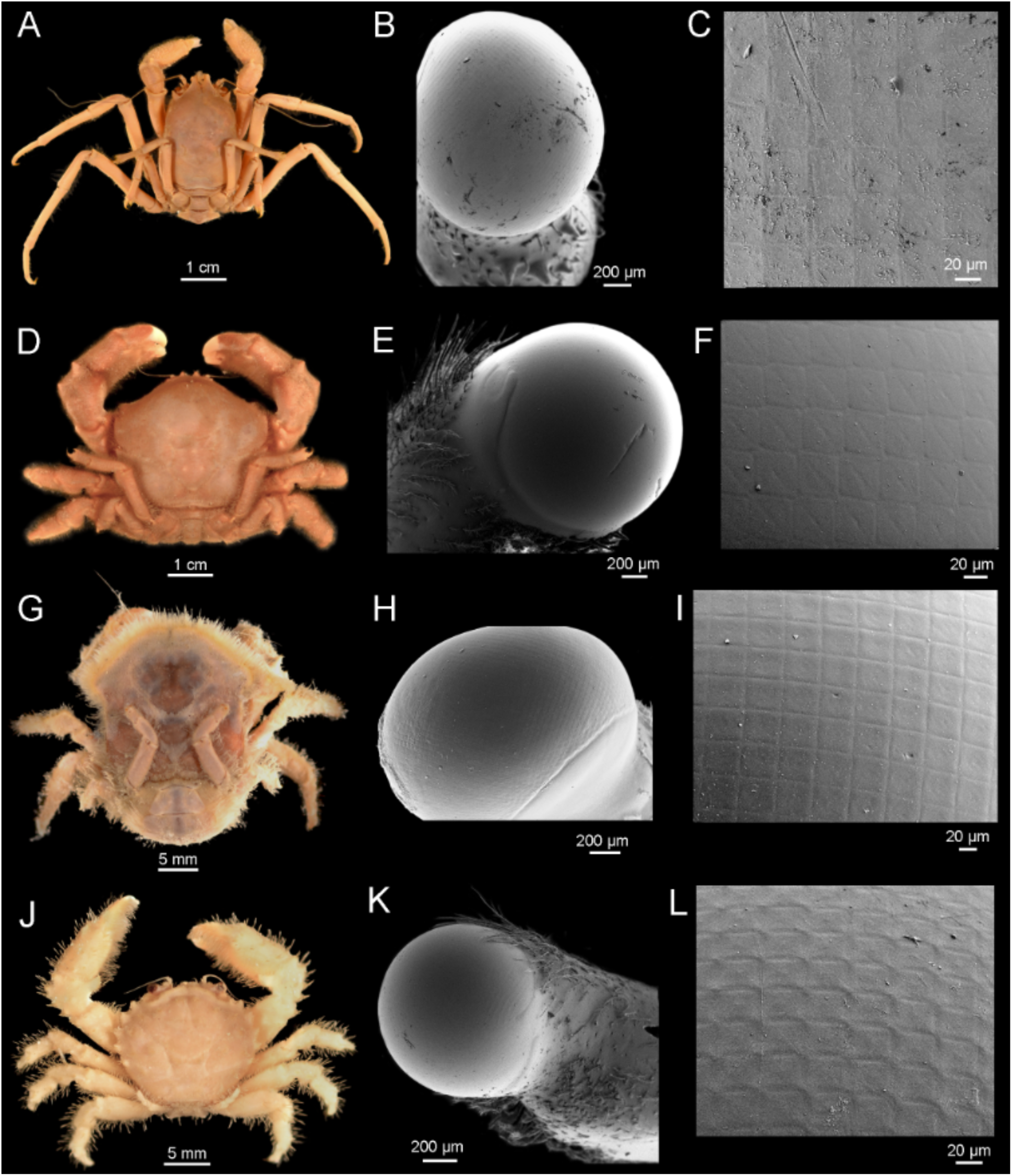
Homolodromioidea and Dromioidea. A–C, Homolodromioidea: Homolodromiidae: *Dicranodromia felderi*, USNM 252207; A, dorsal view of female; B, SEM image of right eye; C, details of the cornea bearing square facets in orthogonal packing. D–I, Dromioidea: Dromiidae; D, ?*Moreiradromia sarraburei*, USNM 1277453, dorsal view of male; E–F, *Dromia personata*, USNM 1277452, female; F, SEM image of right eye; F, details of the cornea bearing square facets in orthogonal packing; G–I, *Hypoconcha* sp., 186466; G, dorsal view of male; H, SEM image of right eye; I, details of the cornea bearing square facets in orthogonal packing. J–L, Dromioidea: Dynomenidae: *Dynomene filholi*, USNM 121402; J, dorsal view of male; K, SEM image of right eye; L, details of the cornea bearing square facets in orthogonal packing. Figure by J. Luque.

**Figure 7.**
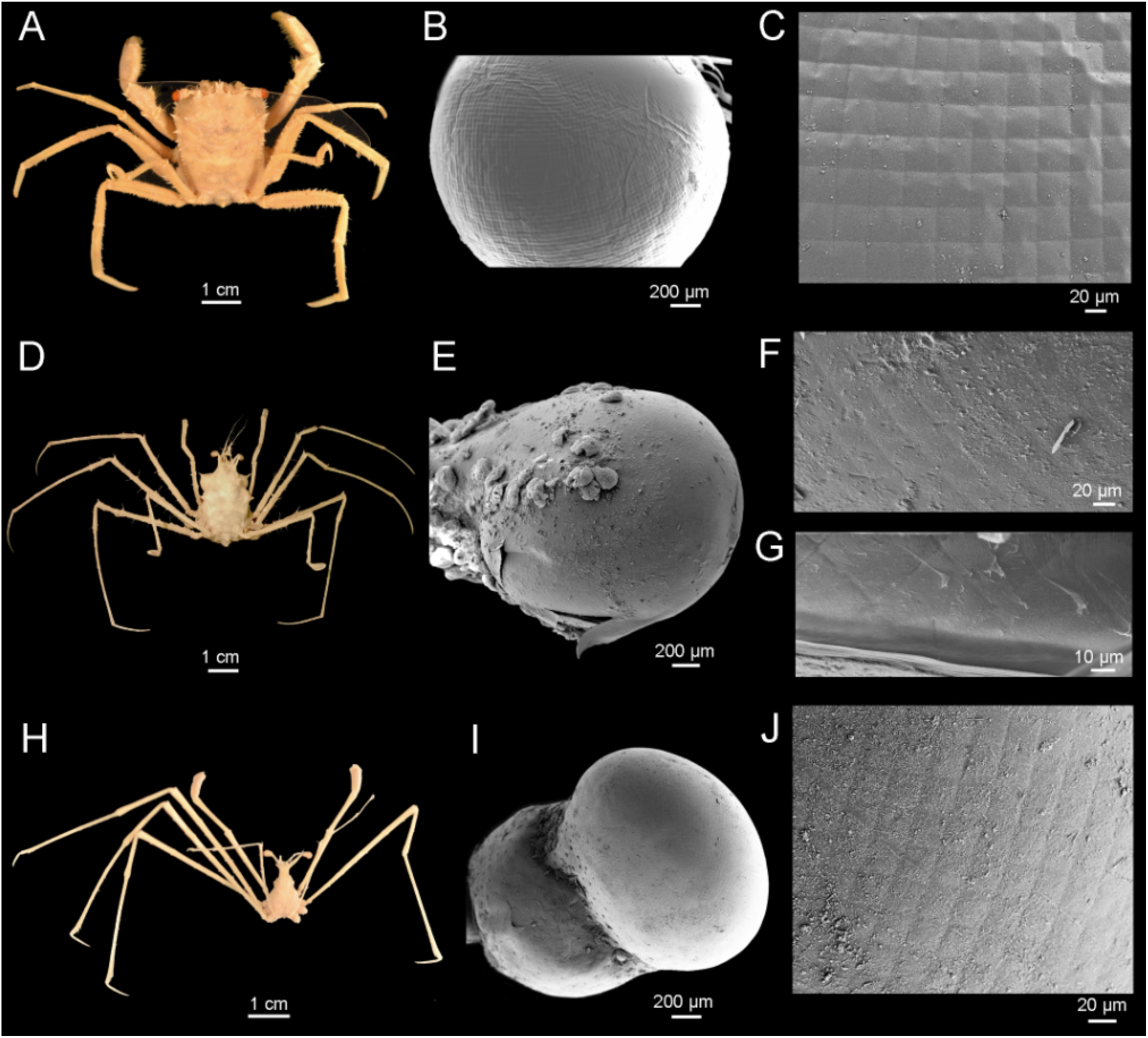
Homoloidea. A–G, Homolidae; A–C, *Homola minima*, USNM 1185786; A, dorsal view of male; B, SEM image of globular right eye; C, details of the cornea bearing square facets in orthogonal packing. D–F, *Latreillopsis bispinosa*, QMW.17070; D, dorsal view; E, SEM image of right eye; F, details of the cornea bearing square facets in orthogonal packing; G, detail of the eye under the cuticle, showing square facets in orthogonal packing. H–I, Latreillidae: *Eplumula phalangium*, USNM 74587; H, dorsal view of male; I, SEM image of right eye and podophthalmite; J, details of the cornea bearing square facets in orthogonal packing. Figure by J. Luque.

**Figure 8.**
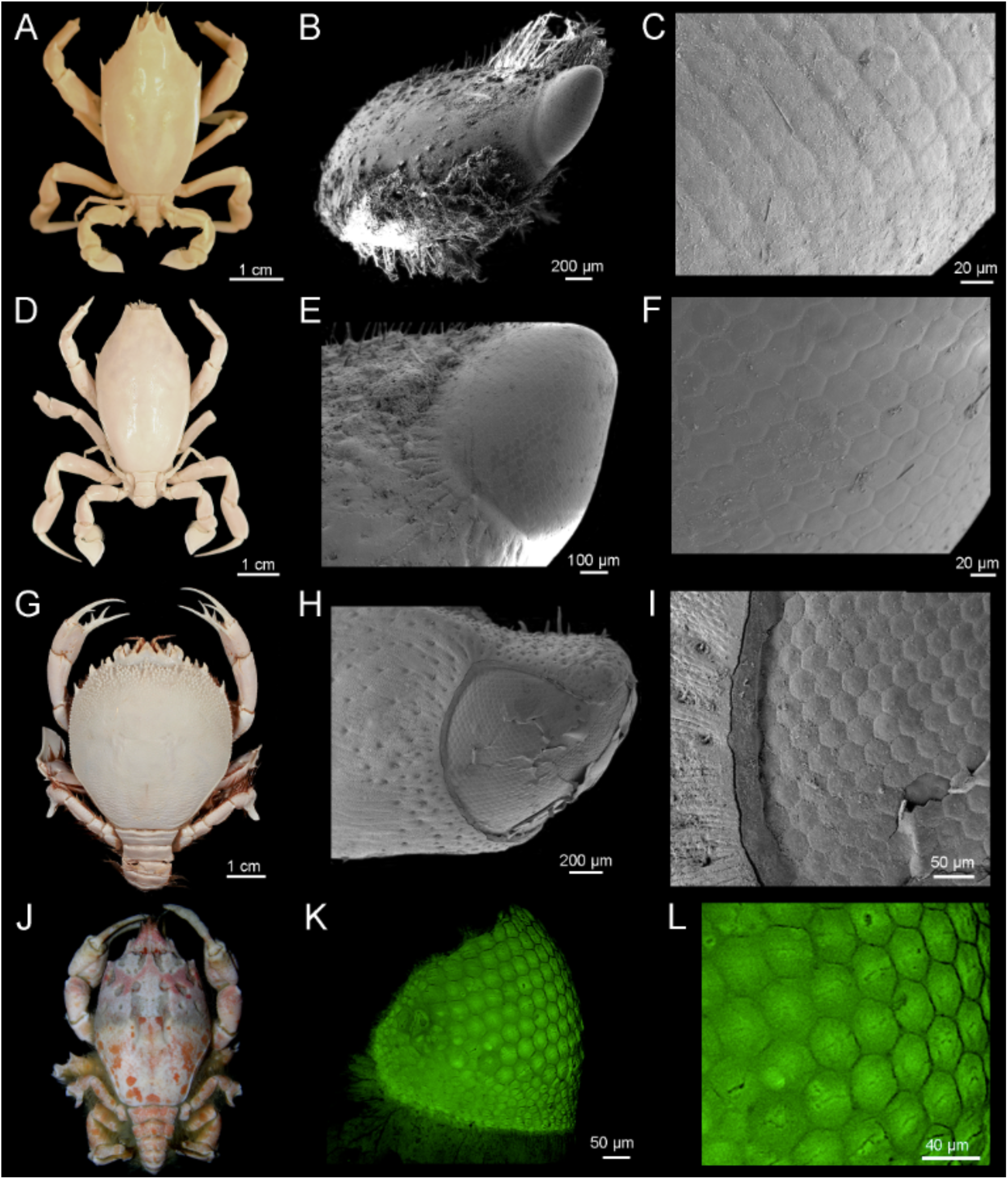
Raninoidea. A–F, Lyreididae; A–C, *Lysirude nitidus* (=*Lyreidus bairdii*), USNM 66638; A, dorsal view of female; B, SEM image of small right eye in a stout podophthalmite; C, details of the cornea bearing hexagonal facets in hexagonal packing. D–F, *Lysirude griffini*, USNM 216726; D, dorsal view of male; E, SEM image of small right eye; C, details of the cornea bearing hexagonal facets in hexagonal packing. G–I, Raninidae: Cyrtorhininae: *Cyrtorhina granulosa*, MNHN-IU-2016-2020 (= MNHN-B16181); G, dorsal view of female; H, SEM image of small right eye; I, details of the cornea bearing hexagonal facets in hexagonal packing. J–L, Raninidae: Symethinae: *Symethis* sp., uncatalogued specimen; J, dorsal view of male; K, Confocal microscope image of small right eye showing the different shapes and sizes of facets through the cornea; L, details of the cornea bearing hexagonal facets in hexagonal packing. Figure by J. Luque.

**Figure 9.**
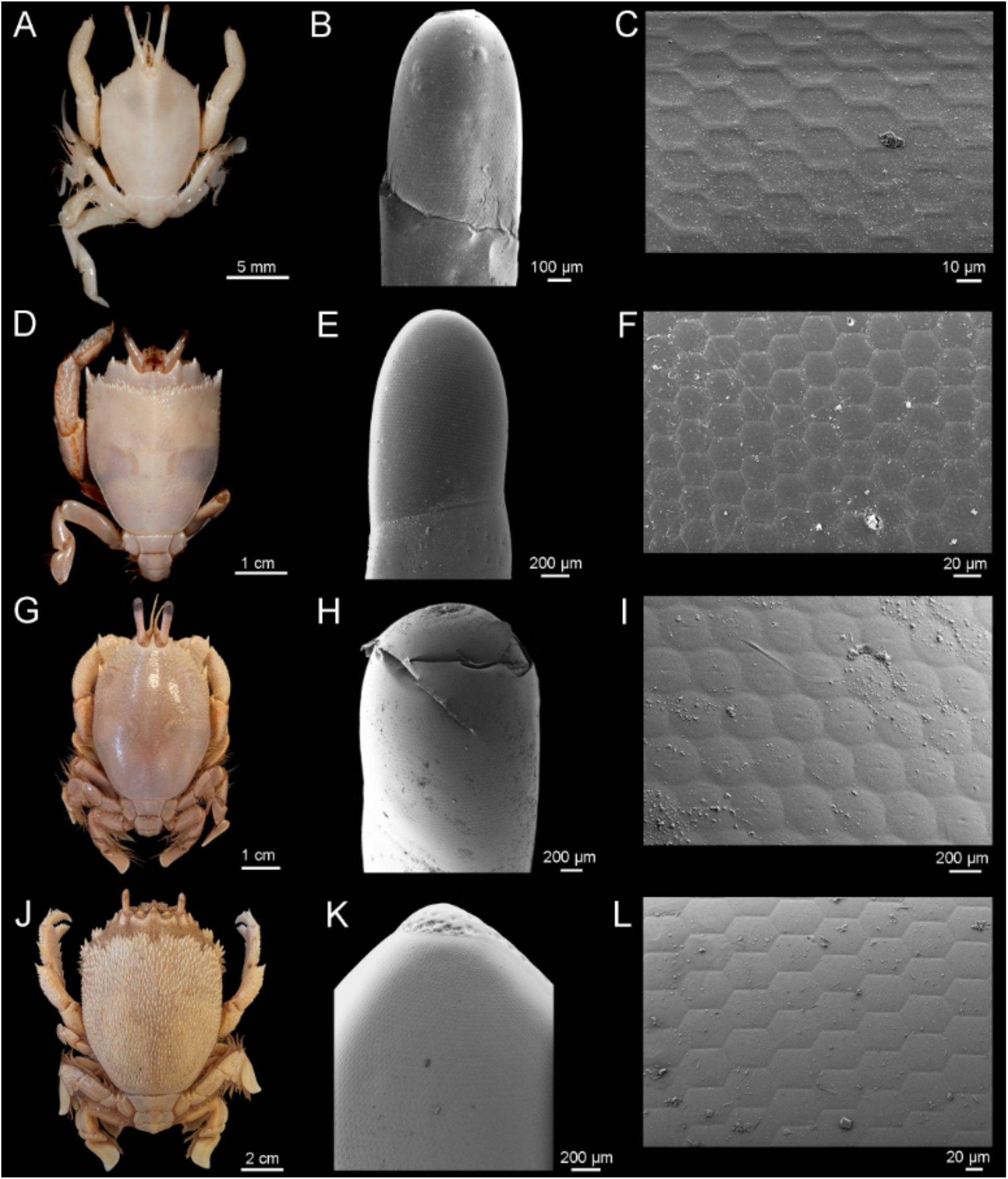
Raninoidea (cont.). A–I, Raninidae: Notopodinae; A–C, *Cosmonotus grayi*, MNHN-IU-2016-2024; A, dorsal view of male; B, SEM image of right eye; C, details of the cornea bearing hexagonal facets in hexagonal packing. D–F, *Notopus dorsipes*, MNHN-IU-2016-2023 (= MNHN-B7933); D, dorsal view of male; E, SEM image of left eye; F, details of the cornea bearing hexagonal facets in hexagonal packing. G–I, *Ranilia muricata*, USNM 121656; G, dorsal view of female; H, SEM image of right eye; I, details of the cornea bearing hexagonal to circular facets in hexagonal packing. J–L, Raninidae: Ranininae: *Ranina ranina*; J, dorsal view of specimen USNM 239219; K–L, specimen USNM 265062, female; K, SEM image of right eye; L, details of the cornea bearing hexagonal facets in hexagonal packing. Figure by J. Luque.

**Figure 10.**
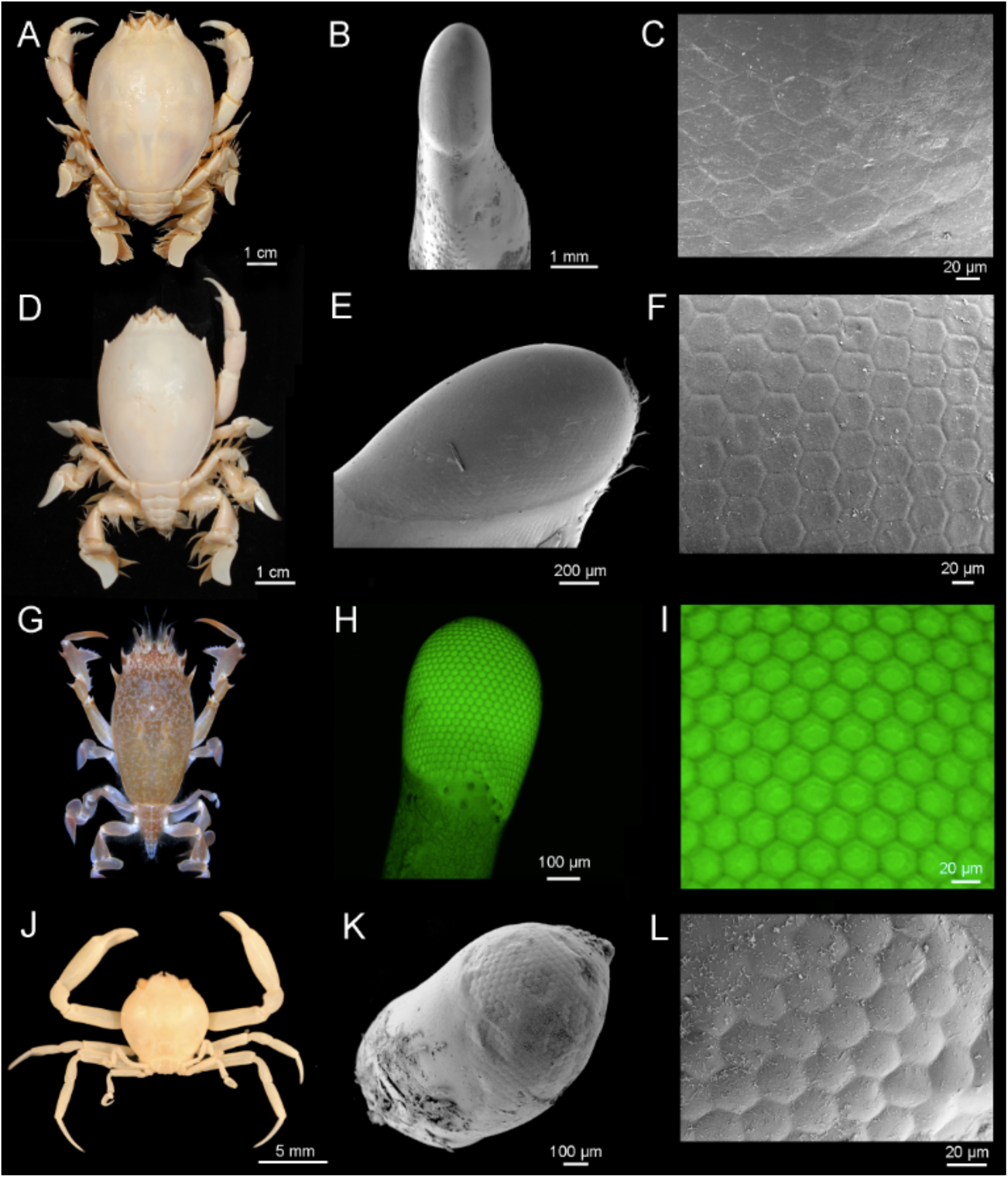
Raninoidea (cont.) and Cyclodorippoidea. A–I, Raninoidea: Raninidae: Raninoidinae: A–C, *Notopoides latus*, MNHN-IU-2016-2025 (= MNHN-B19110); A, dorsal view of male; B, SEM image of right eye and eyestalk; C, details of the cornea bearing hexagonal facets in hexagonal packing. D–F, *Notosceles viaderi*, MNHN-IU-2016-2029 (= MNHN-B28964); D, dorsal view of male; E, SEM image of right eye; F, details of the cornea bearing hexagonal facets in hexagonal packing. G–I, *Raninoides benedicti*, specimen uncatalogued; G, dorsal view of male; H, Confocal microscope image of right eye and eyestalk; I, close up of the cornea bearing hexagonal facets in hexagonal packing. J–L, Cyclodorippoidea: Cyclodorippidae: *Clythrocerus nitidus*, USNM 77380; J, dorsal view of male; B, SEM image of small right eye; C, details of the cornea bearing hexagonal facets in hexagonal packing. Figure by J. Luque. Photo G courtesy of Arthur Anker.

**Table 1.**
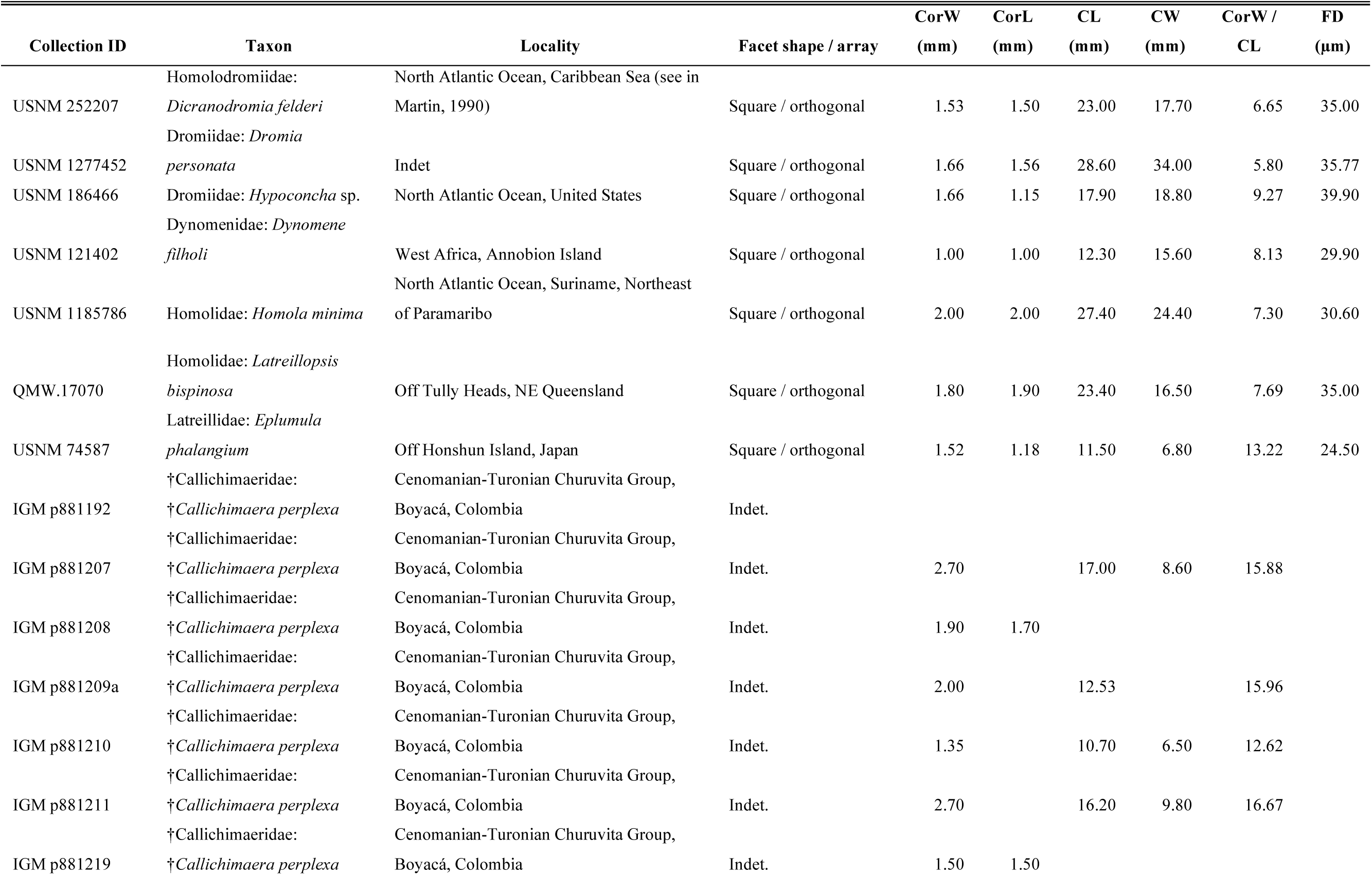
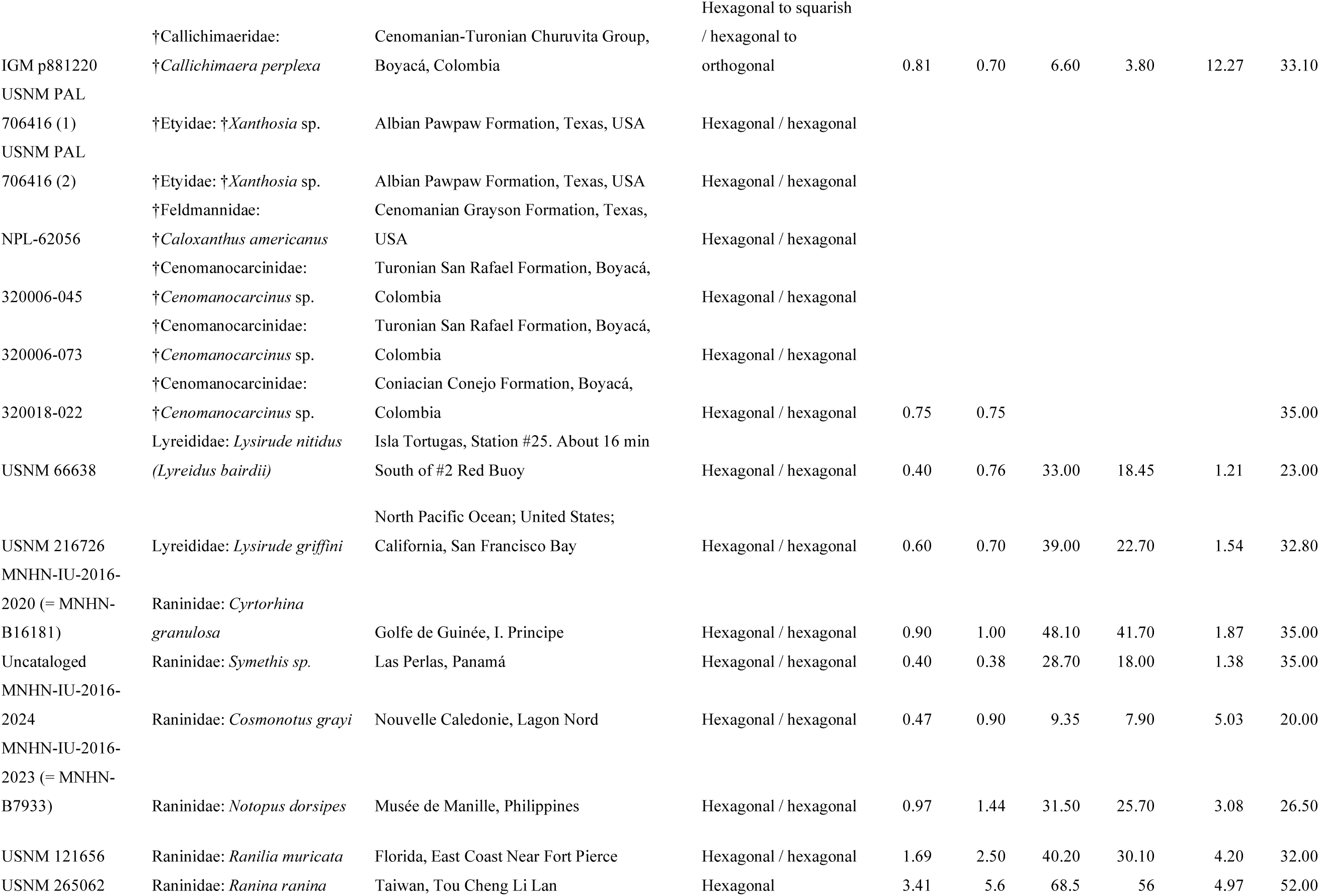
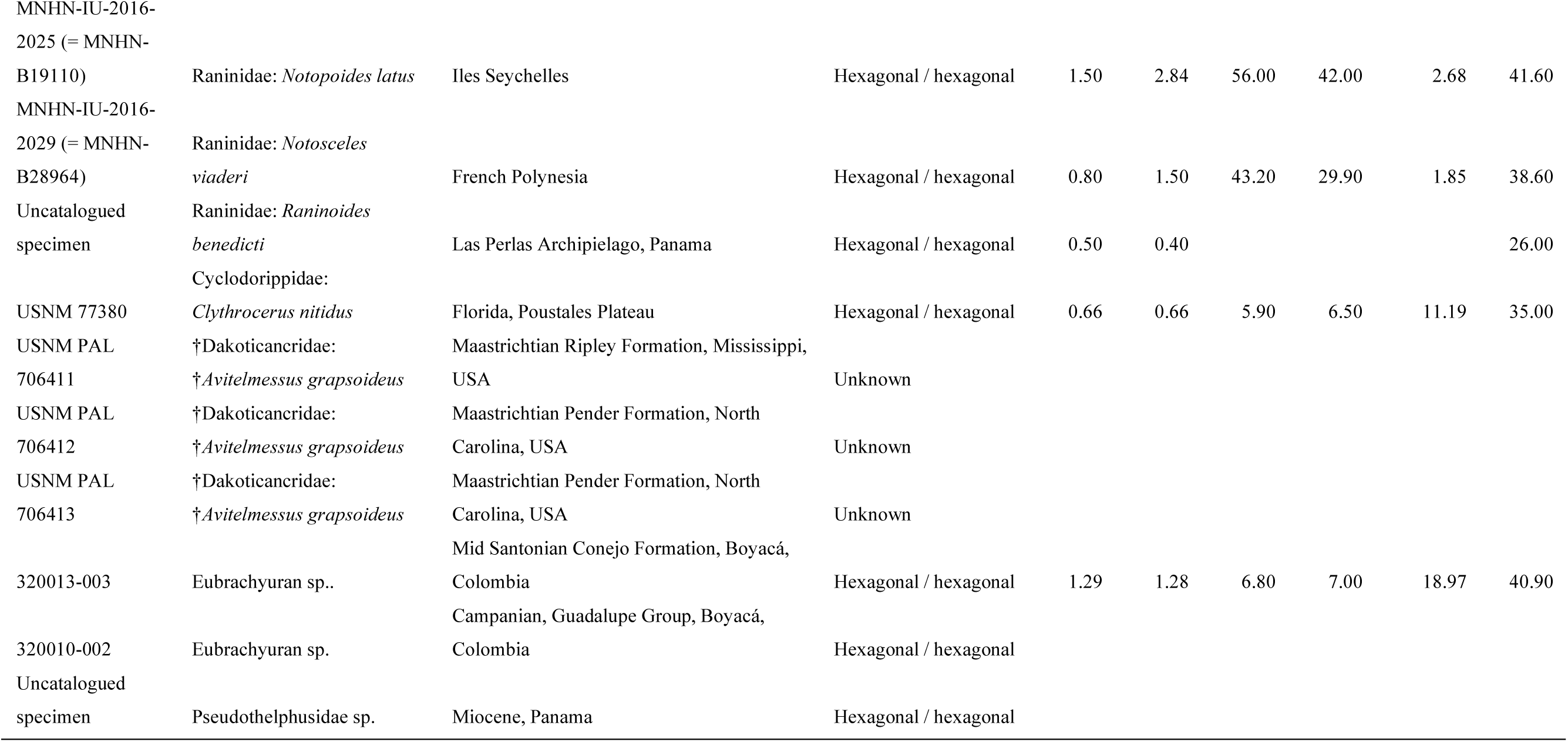
List of extant podotreme and fossil brachyuran specimens studied. Abbreviations: CL: carapace maximum length; CorL: cornea length; CorW: cornea width/diameter; Cw: carapace maximum width; FD: facet diameter; mm: millimeters; µm: microns. Dagger (†) indicates extinct taxa.

#### Fossil taxa

We investigated the ommatidia morphology in eight fossil species of podotreme (five species) and eubrachyuran (three species) crabs with eyes preserved. Studied taxa included †*Callichimaera perplexa* Luque et al., 2019a (†Callichimaeroidea: †Callichimaeridae), from the upper Cenomanian–lower Turonian Churuvita Group (95–90 Mya) of Boyacá, Colombia (Fig. 11); †*Xanthosia* sp. (†Etyoidea: †Etyidae), from the upper Albian Pawpaw Formation of Texas, USA (Fig. 12A–F), and †*Caloxanthus americanus* Rathbun, 1935a (†Etyoidea: †Feldmannidae), from the Cenomanian Grayson Formation of Texas (Fig. 12G–I); †*Cenomanocarcinus* spp. (†Necrocarcinoidea: †Cenomanocarcinidae), from the lower-mid Turonian San Rafael Formation and the upper Coniacian Conejo Formation of Boyacá, Colombia (Fig. 13); †*Avitelmessus grapsoideus* Rathbun, 1935a (†Dakoticancroidea: †Dakoticancridae), from the Maastrichtian Ripley Formation of Mississippi and the Peedee Formation of North Carolina, USA (Fig. 14); two species of fossil Eubrachyura, from lower-mid Santonian Conejo Formation and the Campanian of Boyacá, Colombia (Fig. 15A–F); and a fossil freshwater crab from the lower Miocene Pedro Miguel Formation of the Panama Canal expansion zone, Panama (Fig. 15G–I). A complete list of the fossils studied and their associated information is provided in Table 1.

**Figure 11.**
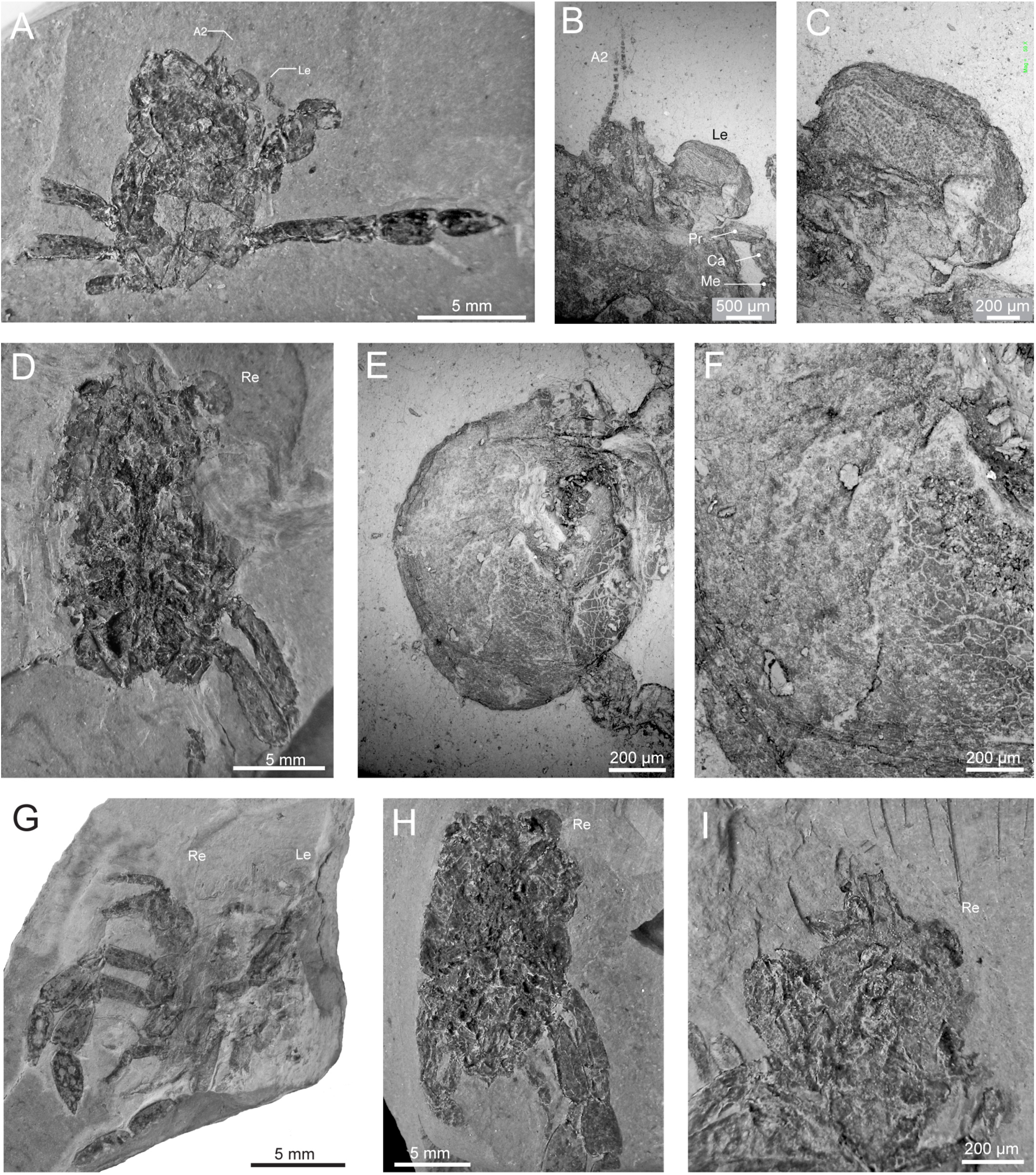
Specimens of the crab †*Callichimaera perplexa* (†Callichimaeroidea: †Callichimaeridae) with eyes preserved. Specimens coated with ammonium chloride, except for SEM images. A–C, Paratype IGM p881210, ventral view; A, specimen showing the second antennae and left compound eye; B, SEM of anterior portion, showing the mxp3, antennae, and left compound eye; C, SEM image showing details of the facets. D–F: Paratype IGM p881207; D, specimen showing legs P2–P3, and right eye; E, SEM image of right eye; F, SEM close–up of the same eye, showing facets in hexagonal arrangement. G, Paratype IGM p881219, ventral view showing the chelipeds, legs P2–P5, both eyes, and rostrum. H, Paratype IGM p881211, showing right eye. I, Paratype IGM p881192, showing a preserved eye. Abbreviations: A2: second antenna (antenna s.s.); Ca: carpus; Le–Re: left and right eyes; Me: merus; Pr: propodus. A, D, G–I coated with ammonium chloride; B–C, E–F, dry and uncoated SEM images. Figure by J. Luque.

**Figure 12.**
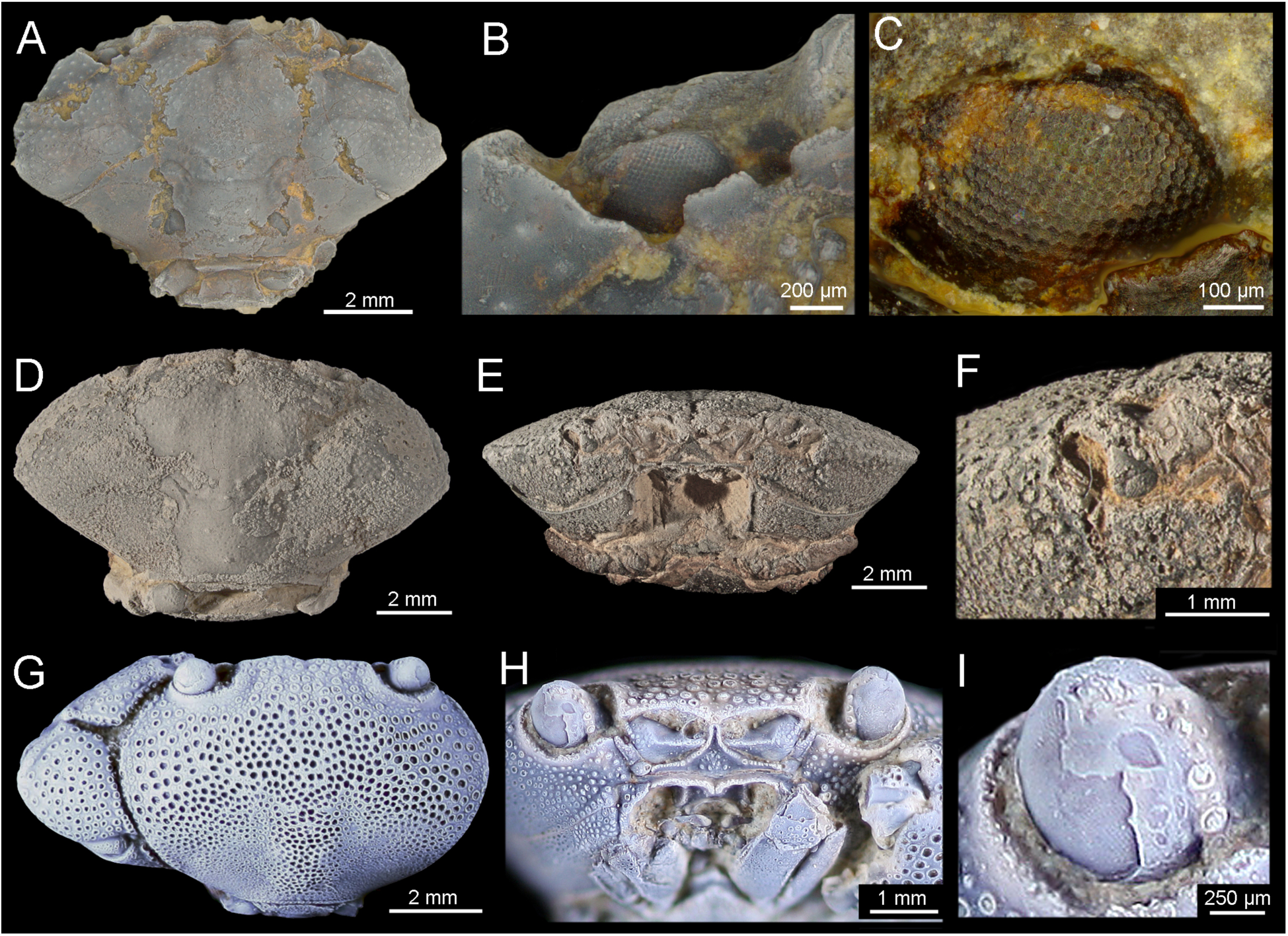
Specimens of the fossil crabs †*Xanthosia* (†Etyoidea: †Etyidae) and †*Caloxanthus* (†Etyoidea: †Feldmannidae) with eyes preserved. A–C, †*Xanthosia* sp., specimen USNM PAL 706416 (1); A, dorsal view; B, close-up of dorsal left eye and orbital margin; C, close-up of left compound eye bearing hexagonal facets in hexagonal array. D–F, †*Xanthosia* sp., specimen USNM PAL 706416 (2); D, dorsal view; E, frontal view showing the orbits; F, close-up of right eye with fragmented eye cornea bearing hexagonal facets in hexagonal packing. G–I, †*Caloxanthus americanus*, specimen NPL-62056; G, dorsal view; H, close-up of frontal view showing the hemispherical eyes; I, close-up of right compound eye bearing hexagonal facets in hexagonal array. All specimens photographed dry and coated with ammonium chloride, except for C (uncoated). Dagger (†) indicates extinct taxa. Images G–I courtesy of Liath Appleton (University of Texas). Figure by J. Luque.

**Figure 13.**
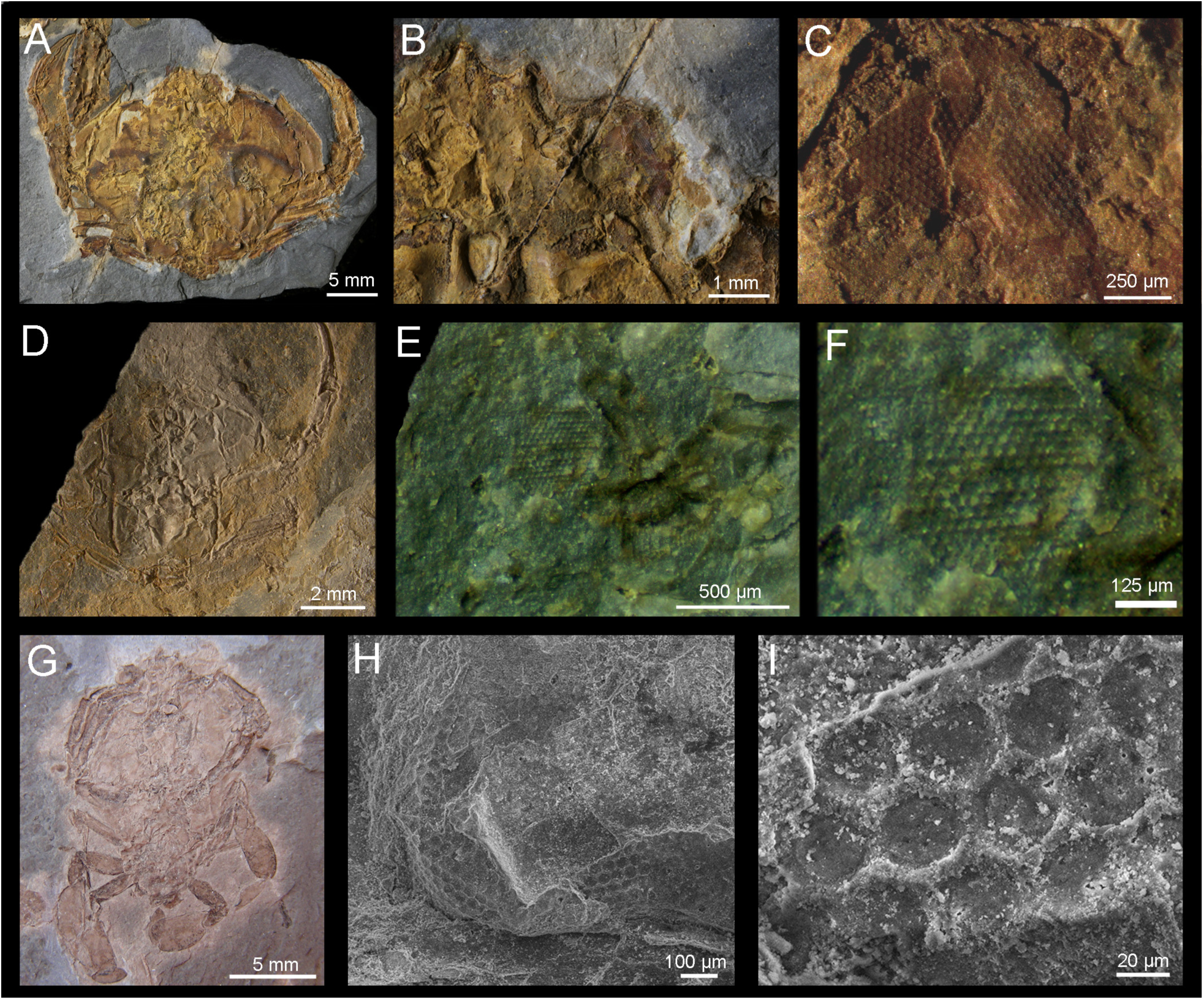
Specimens of the crab †*Cenomanocarcinus* (Raninoida: †Necrocarcinoidea: †Cenomanocarcinidae) with eyes preserved. A–C, Specimen 32006-073, San Rafael Formation, lower Upper Cretaceous (Turonian, ∼90 Ma), Boyacá, Colombia; A, dorsal view; B, close-up of right eye; C, close-up of right eye cornea bearing hexagonal facets in hexagonal packing. D–F, Specimen 320006-045, San Rafael Formation, lower Upper Cretaceous (Turonian, ∼90 Ma), Boyacá, Colombia; D, dorsal view; E, close-up of left eye; F, close-up of left eye cornea bearing hexagonal facets in hexagonal packing. G–I, Specimen 320018–022, Conejo Formation, Upper Cretaceous (Coniacian, 85 Ma) of Boyacá, Colombia; G, ventral view of female; H, SEM image of right eye (Re); I, close-up of ventral right eye cornea showing small hexagonal facets in hexagonal packing. All specimens photographed dry, uncoated. Dagger (†) indicates extinct taxa. Figure by J. Luque.

**Figure 14.**
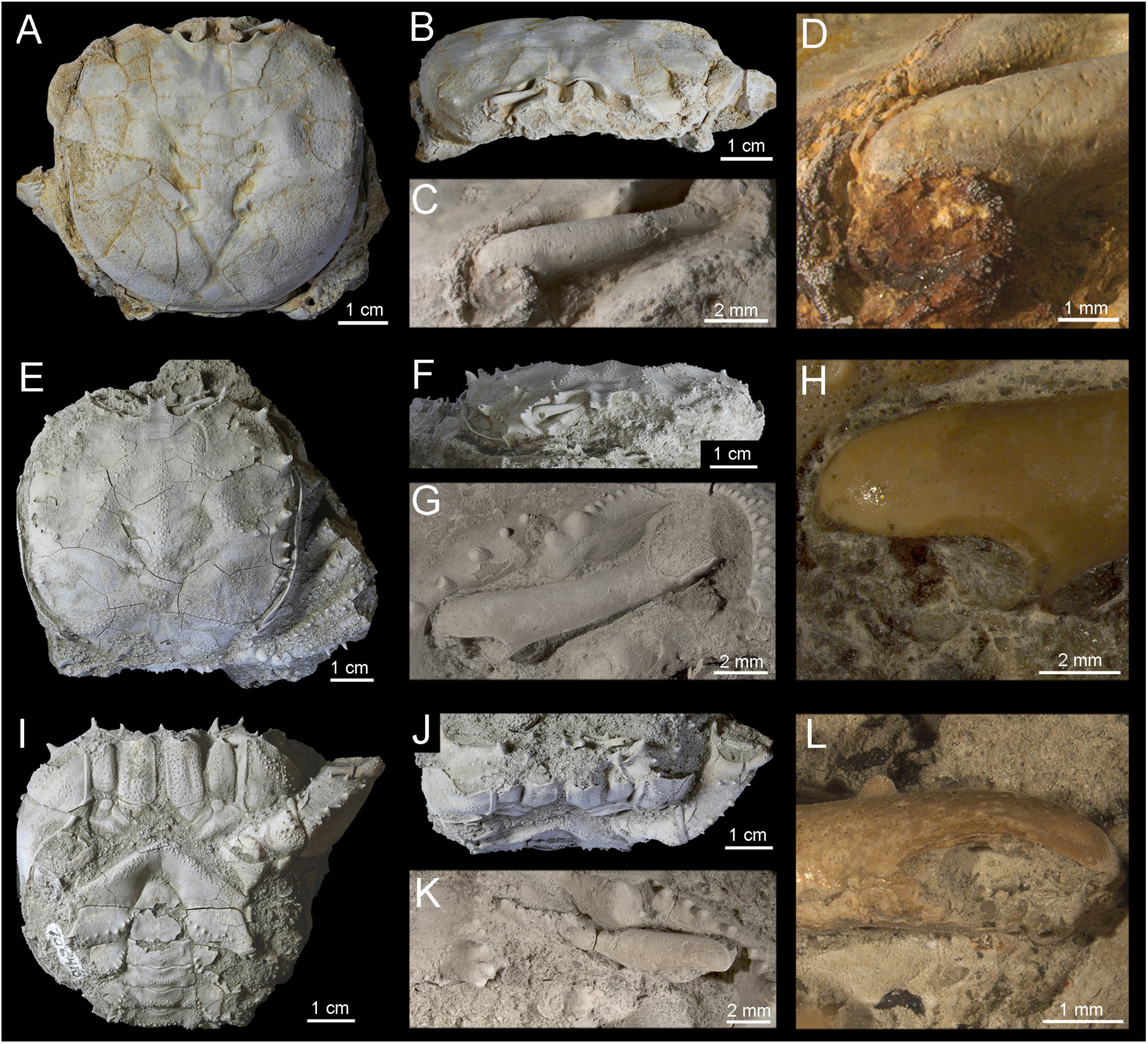
Specimens of the crab †*Avitelmessus grapsoideus* (†Dakoticancroidea: †Dakoticancridae) with eyes preserved. A–D, USNM PAL 706411, Ripley Formation, Upper Cretaceous (Maastrichtian), Mississippi, USA; A, dorsal view; B, frontal view; C, close-up of right eye and eyestalk; D, close-up of distal eye, cornea missing. E–H, USNM PAL 706412, Pender Formation, Upper Cretaceous (Maastrichtian), North Carolina, USA; E, dorsal view; F, frontal view; G, close-up of right eye and eyestalk; H, close-up of distal eye, cornea missing. I– L, USNM PAL 706413, Pender Formation, Upper Cretaceous (Maastrichtian), North Carolina, USA; I, ventral view; J, frontal view; K, close-up of left eye and eyestalk; L, close-up of distal eye, cornea missing. All specimens photographed dry; A–C, E–G, I–K coated with ammonium chloride; D, H, L uncoated. Dagger (†) indicates extinct taxa. Figure by J. Luque.

**Figure 15.**
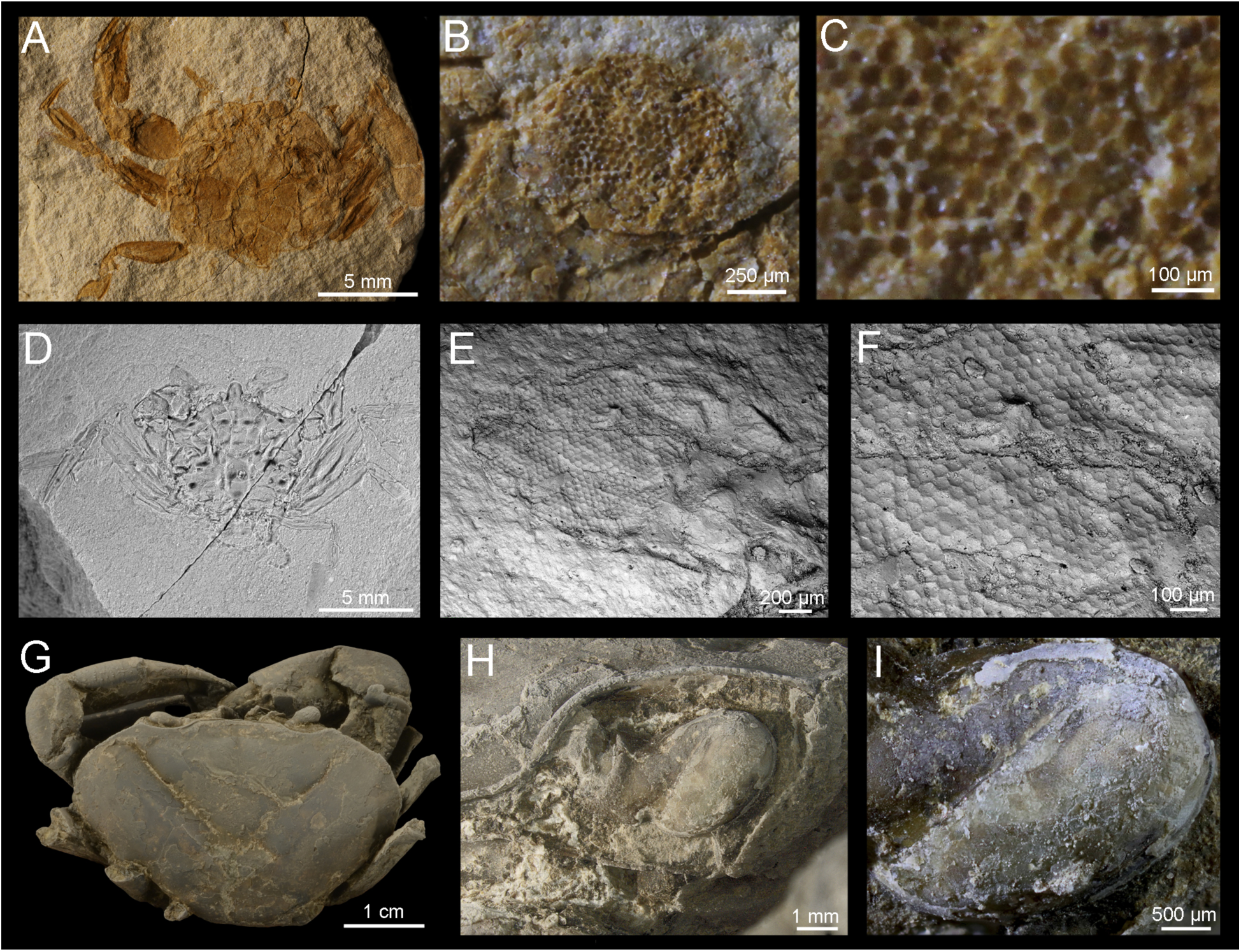
Some fossil Eubrachyura crabs preserving compound eyes. A–C, Eubrachyura sp., Campanian (Upper Cretaceous, ∼80–75 Mya) of Boyacá, Colombia; A, ventral view of specimen IGM p, 320010-002; B, close-up of eye and orbit; C, close-up of cornea bearing small hexagonal facets packed hexagonally. D–F: Eubrachyura sp., lower-mid Santonian (Upper Cretaceous, ∼85 Mya) of Boyacá, Colombia; D, negative of male dorsal carapace showing the pereiopods, pleon, chelipeds, and large compound eyes; E, SEM image of compound eye bearing facets; F, close-up showing small hexagonal facets in hexagonal packing. G–I, Pseudothelphusoidea: Pseudothelphusidae indet., uncatalogued specimen, Miocene of Panama, Panama; G, frontal view showing the fronto-orbital region, the 3^rd^ maxillipeds, and the compound eyes; H, close-up of the left eye; I, details of the cornea preserving hexagonal facets in hexagonal packing, although hardly discernible. Dagger (†) indicates extinct taxa. Figure by J. Luque.

### Methods

#### Tissue processing

Eyes of selected extant adult crabs from museum collections, preserved in 70% EtOH were dissected and prepared for Scanning Electron Microscope (SEM) via dehydration through a series of rinses in EtOH at 70%, 90%, and twice at 100% at intervals of 20 and 30 minutes for small and large samples, respectively. Then the tissues were rinsed for similar time intervals in a mixture of EtOH and Hexamethyldisilazane (HMDS) at 25:75, 50:50, and 75:25 ratios, plus two final rinses in 100% HDMS. This tissue dehydration technique is faster, easier, and less expensive than the critical point drying with CO2.

#### Imaging

Most fossils were coated with sublimated NH_4_Cl prior to photographing whole specimens to enhance relief and fine ornament. Sets of photographs at different focal points were taken with a Nikon Eclipse 80i + Nikon Digital Camera Dxm 1200f, Olympus SZX16® Research Stereomicroscope with a digital camera Qimaging Retiga 2000R Fast 1394, and a Leica Macroscope with Spotflex digital camera. The resulting multi-layered stacks of photos were merged in a single high–definition image using the stacking software Helicon Focus stacking software. Extant specimens were photographed with a Nikon Digital Camera D3100 with MicroNikkor 60 mm and 105 mm lenses.

Dissected and mounted eyes from fossil and extant crabs were studied under Zeiss Scanning Electron Microscope (SEM) Evo 40vp under low vacuum and variable pressure and Back-scattered Electron Detector (BSED) with acceleration voltages of 15 and 20kV, and under a Zeiss Sigma 300 VP-FESEM scanning electron microscope at the Smithsonian Tropical Research Institute, Panama (STRI), and the University of Alberta, Edmonton, Canada. All eye samples from extant taxa were coated with Au/Pd prior to SEM imaging, except from two specimens imaged using an Olympus FV1000 Confocal Microscope.

#### Lens packing and facet measurements

A strongly supported phylogenetic framework is missing for podotremes (Fig. 16), and is the subject of the present investigation, therefore to assess the sources of variation in lens shape and packing among brachyuran crabs, we utilized geometric morphometrics (Bookstein, 1991). Simple measurements of cornea dimensions and facet diameters (Table 1) were obtained using ImageJ. For morphometric measurements, we examined seventeen museum specimens representative of sixteen genera, all representing adult individuals (Table 1). For each specimen, we determined whether square or hexagonal facets were present. Lens packing was represented by four two-dimensional landmarks taken at the centroid of four adjacent lenses (Fig. 17A,B). From each individual, we collected five sets of landmarks from different areas of the eye, except in cases where the curvature of the eye or damage to the ommatidia severely affected landmark acquisition. All landmark measurements were taken from high magnification images (Figs 6–15), and measured using the software TpsDig (Rohlf, 2005), which translates landmarks into Cartesian coordinates. Those Cartesian coordinates, as transformed landmarks, underwent Procrustes superimposition in PAST (Hammer et al., 2001) to remove the effects of rotation, translation, and size, which minimizes the shape difference between sets of landmarks (Bookstein, 1991). This procedure translates all specimens to the origin, scales them, and rotates them to minimize deviations of landmark coordinates to an average configuration for all specimens. Procrustes superimposition also transforms the Cartesian coordinates originally taken during landmark acquisition into Procrustes coordinates (i.e., this creates a covariance matrix of Procrustes residuals). Because of this, axes are scaled in Procrustes units which are arbitrary units that only serve as a within-study metric. Principal components (PCs) were calculated from the covariance matrix of the Procrustes coordinates (Dryden and Mardia, 1998), and that principal components space is often referred to as a ‘morphospace’ (Fig 17C). Each data point in that morphospace corresponds to a particular shape configuration. The closer two data points are, the more similar they are in shape. In a principle components analysis of landmark data, each PC describes a different component of shape, and PC axes are uncorrelated.

**Figure 16.**
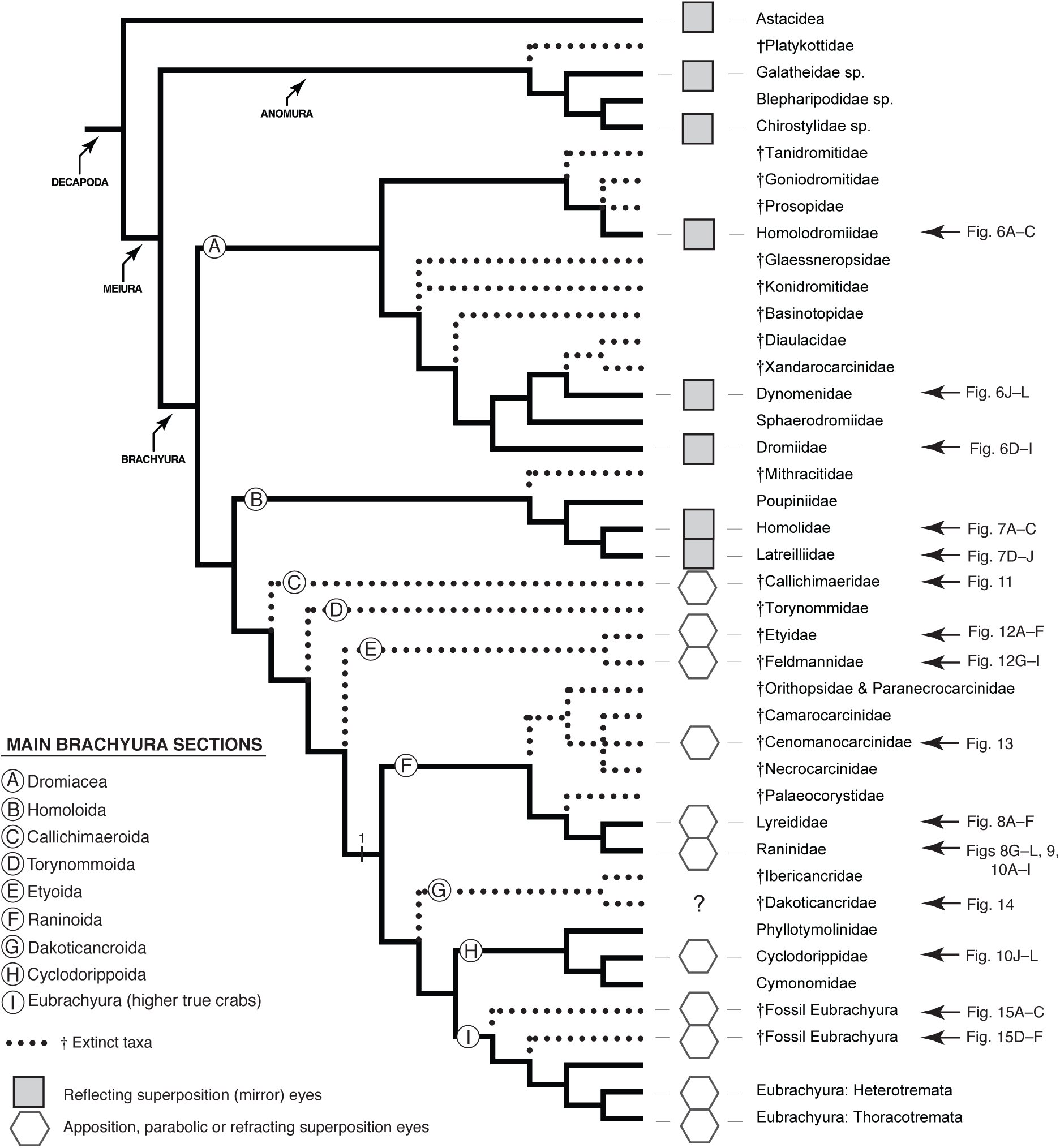
Distribution of visual systems in Brachyura. Hexagons: hexagonal facets with hexagonal packing, typical of apposition, parabolic superposition, and refracting superposition eyes. Squares: square facets in an orthogonal array typical of reflecting superposition eyes, or ‘mirror’ eyes. Only the eye facet shape of taxa examined in this studied are indicated. Figure by J. Luque. Tree topology modified after Luque et al. (2019a).

**Figure 17.**
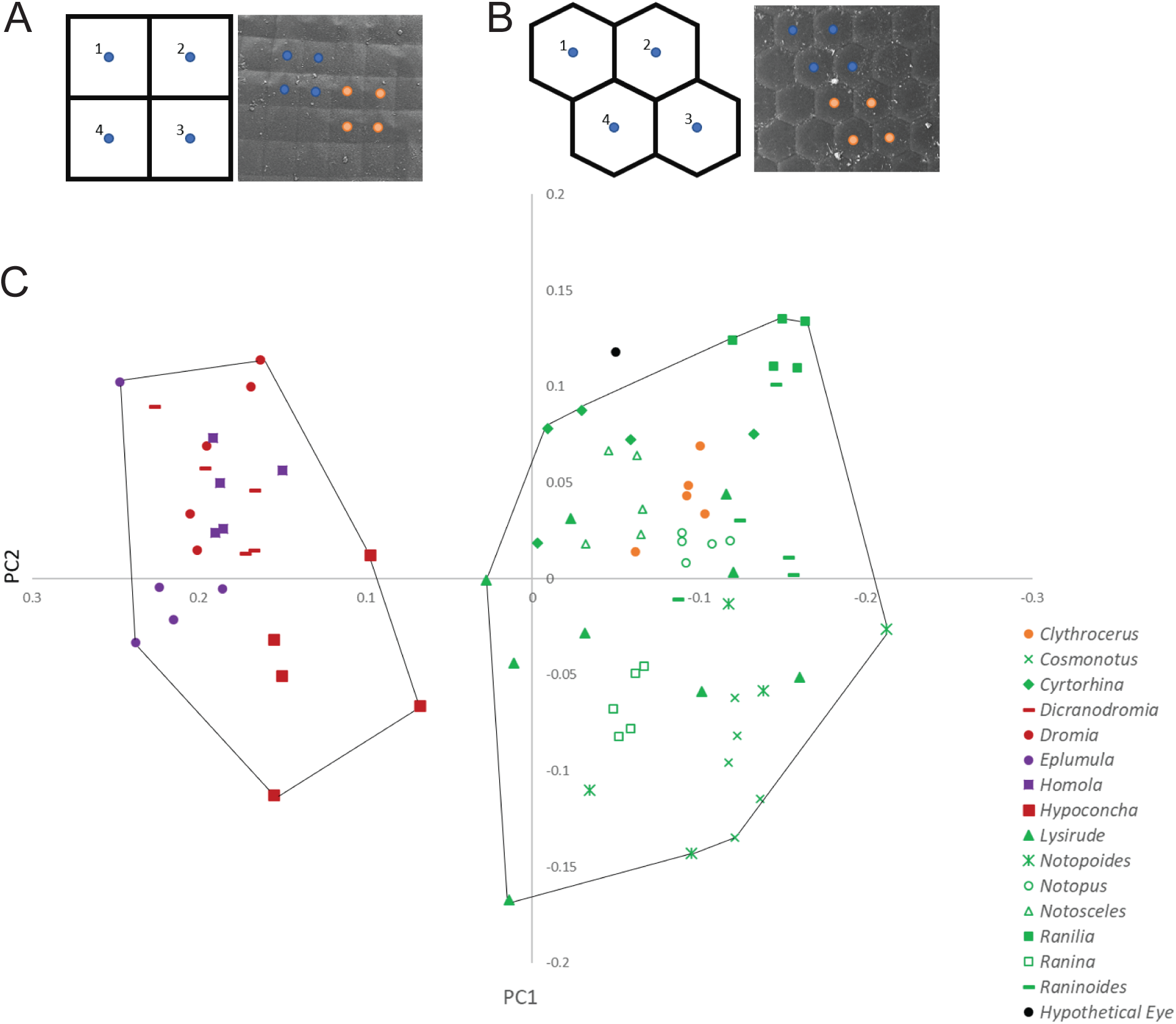
Principle component analysis resulting from geometric morphometric analysis of lens packing. A, Location of landmarks taken for geometric morphometric analysis in both orthogonal/square and hexagonally packed eyes. B, Results from GMM analysis. Two distinct groupings of eye types are revealed through GMM analysis, an orthogonal packing arrangement of square facets, and an hexagonally packed arrangement of hexagonal to roundish facets. Taxa are further grouped by facet shape (also orthogonal vs. hexagonal), showing that there is no overlap in morphology between lens packing and facet shape. Podotreme crab taxa with orthogonally arranged facets are represented by red and purple icons, and taxa with hexagonally arranged facets are represented by green and orange icons. Figure by K.M. Jenkins.

To assess the relationship between facet shape and packing by geometric morphometrics (as outlined above), we performed a multivariate analysis of variance (MANOVA) from PC scores in PAST (Hammer et al., 2001). Facet shape was treated as a categorical variable (i.e., hexagonal vs. square).

## RESULTS

Below, we describe the gross, facet, and packing morphology recorded for the eyes of fossil and extant species of podotreme crabs here investigated.

### Eyes of extant podotreme brachyurans

#### Homolodromioidea

*Dicranodromia felderi* has globular eyes slightly larger than the eyestalk. The podophthalmite is partly covered dorsally in small fine to conical spines, and the eye and eyestalk partially fit a shallow orbit laterally bounded by a short, triangular, anterolaterally diverging outer orbital spine (Fig. 6A). In the studied specimen, the corneal eye is nearly as wide as long, its width is approximately 6% the carapace, and is covered in small ommatidia (35 µm in diameter) with square facets packed in a rectilinear lattice (Fig. 6B,C; Table 1).

#### Dromioidea

The dromiids *Dromia personata* (Fig. 6E,F), *Hypoconcha* sp. (Fig. 6G–I), and the dynomenid *Dynomene filholi* (Fig. 6J–L) all have eyes with square facets in an orthogonal array. In *Dromia personata*, the eye is small, globular, and about as long as the eyestalk. The podophthalmite is covered with plumose setae, where secondary acicular setae stem from the primary setae (Fig. 6E). Its corneal surface is nearly as wide as long, with a width diameter less than 6% the carapace length, and is covered in small square ommatidia (35 µm diameter) (Fig. 6F; Table 1). In *Hypoconcha* sp., the eyes are also globular, wider than long, and slightly longer than the eyestalk. The cornea has a diameter that is about 9% of the carapace length, and is covered with small rhomboid ommatidia (40 µm) (Fig. 6H,I). In *Dynomene filholi*, the eyes are small and globular, and shorter than the eyestalk, and like in *Dromia*, the podophthalmite is covered with plumose setae (Fig. 6K). The cornea has a diameter about 8% the carapace length, and it is covered with small square ommatidia (40 µm in diameter) with depressed edges (Fig. 6L).

Square facets packed in an orthogonal lattice have been previously reported for other dromiids and dynomenids such as *Dromia vulgaris* and *Dynomene pilumnoides* (Gaten, 1998; Scholtz and McLay, 2009; D. Guinot, pers. comm. to JL, 2016), supporting the distribution of these features across genera of Homolodromioidea and Dromioidea crabs.

#### Homoloidea

The homolid crab *Homola minima* has hemispherical globular eyes that rest on a cylindrical podophthalmite that is slightly longer than the corneal eye. The basophthalmite is slender, cylindrical, and more than twice as long as the corneal eye or the podophthalmite (Fig. 7A). As in other homolids, the podophthalmite rests on a depressed space acting as a false orbit (Davie et al., 2015). In the studied specimen, the corneal eye is nearly as wide as long, its width is approximately 7% the carapace length, and is covered with small square ommatidia (approx. 30 µm in diameter) packed in an orthogonal lattice (Fig. 7B,C; Table 1). Similarly, the homolid *Latreillopsis bispinosa* has a large globular eye with a short podophthalmite and a slender and much longer basophthalmite (Fig. 7D). The cornea is smooth, nearly as wide as long, its width is less than 8% the carapace length, and is covered in small square ommatidia (35 µm) packed in an orthogonal lattice (Fig. 7F; Table 1). The boundaries between facets are less conspicuous than in the eye of *Homola minima*, but the overall facet shape and array is still evident above and below the cuticle (Fig. 7F,G). The latreillid *Eplumula phalangium* has globular eyes, but they rest in shorter podophthalmites compared to the other homoloid species studied. The cornea is wider than it is long, about 13% as wide as the carapace maximum length, and is covered with small square facets (24.5 µm in diameter) in orthogonal array.

Similar facet shapes and packing match previous findings for other homoloid taxa such as *Paromola cuvieri*, for which eyes of the reflecting superposition (mirror) type have been reported (Gaten, 1998).

#### Raninoidea

The lyreidid crab *Lyreidus nitidus* (Fig. 8A) has small sub-conical eyes resting in stout and much longer podophthalmite, nearly 66% larger than the corneal eye, and covered in fibrous setae (Fig. 8B). The eye and eyestalk are partially protected by a narrow orbit with one supraorbital fissure, and a produced, acute, triangular outer orbital spine directed anteromesially. The cornea width is approximately 1.2% the length of the carapace, and it is constituted by a few hundred small hexagonal facets that are packed in hexagonal array, with an average facet diameter of 23 µm (Fig. 8C; Table 1). The studied specimen of *Lysirude griffini* (Fig. 8D) has even more reduced sub-conical eyes and a longer and broader podophthalmite than *L. nitidus*; the podophthalmite is nearly twice as long as the corneal eye, and it is partly covered in small setae (Fig. 8E). Both eye and eyestalk are barely protected by a narrow orbit with one supraorbital fissure, and a short, blunt, triangular outer orbital spine directed anteriorly (Fig 8D). The cornea width is approximately 1.5% the length of the carapace, and it is constituted by a few hundred small hexagonal facets that are packed in hexagonal array, with an average facet diameter of 23 µm (Fig. 8C; Table 1). Hexagonal facets in hexagonal array have been reported for *Lyreidus tridentatus* (see Scholtz and McLay, 2009), suggesting a shared lack of ‘mirror’ eyes among crabs of the family Lyreididae.

Among all the podotreme crabs studied, those of the family Raninidae have the broadest range of eye shapes, sizes, and orbital constructions. In the subfamily Cyrtorhininae, *Cyrtorhina granulosa* (Fig. 8G) shows a considerable reduction of the corneal region compared with the rest of the eyestalk. Its cornea is sub-conical and dorsally truncated by an extension of the cuticle of the podophthalmite that extends towards the pole of the eye, further reducing the area occupied by the cornea (Fig. 8H). The cornea width is approximately 1.8% the length of the carapace, and it is constituted by small hexagonal facets in hexagonal array, with an average facet diameter of 35 µm (Fig. 8H, I; Table 1). The facets across the cornea are similar in size. The cuticular lenses in *C. granulosa* include a thin epicuticle forming the slightly convex outer facets, an underlying thin exocuticle with a concave center, and a membranous underlying endocuticle forming concave facets with raised edges. The podophthalmite is three times larger than the cornea. It is covered in microcuticular tuberculations and bears multiple setal pits nucleated by a single reduced seta in each. The medial and proximal dorsal portions of the podophthalmite are ornamented with several sub-conical to fungiform nodes ranging in size, the largest of which are capped by an eroded roundish top. Short orbits barely protect the eyes, with a sub-horizontal supraorbital margin bearing two fissures separating the short, blunt, triangular orbital spines.

Species of the subfamily Symethinae have the shortest eyes of all the raninoids studied. In *Symethis* sp. (Fig. 8J) the eyestalk is very reduced, and the corneal eye is concealed in a very narrow orbit, considerably restricting the motion of the eye. The cornea maximum width is about 1.4% of carapace length. The facets across the cornea are quite different in shape and size; the most central facets are hexagonal in hexagonal packing (about 35 µm), while the peripheral facets towards the eyestalk are considerably smaller (around 14.5 µm) and with irregular shapes and packing (Fig. 8K, L; Table 1).

The genera in the subfamily Notopodinae all have well developed eyes on long eyestalks. In *Cosmonotus grayi*, the length of the podophthalmite seems to be the most extreme across raninoids, measuring half the length of the dorsal carapace (Fig. 9A). Its cornea is sub-cylindrical (Fig. 9B), longer than wide, approximately 5% the length of the carapace, and it bears small flattened hexagonal facets around 20 µm in diameter, with hexagonal packing, and with raised facet edges (Fig. 9C; Table 1). *Notopus dorsipes* (Fig. 9D) and *Ranilia muricata* (Fig. 9G) also have corneae that are longer than wide, sub-cylindrical, with a diameter about 3% the carapace length, and three times shorter than the eyestalk. Their facets are also hexagonal to roundish, well defined, and packed in a hexagonal array. The facet diameter in *N. dorsipes* is around 26.5 µm (Fig. 9E,F), and 32 µm in *R. muricata* (Fig. 9H,I; Table 1).

The subfamily Ranininae has only one living genus and species, *Ranina ranina*. It is the largest of all raninoids, reaching carapace length sizes over 15 cm (Luque, unpublished data). *Ranina* eyes are elongate, elliptical to sub-cylindrical (Fig. 9J,K). The eyestalk has a long podophthalmite twice as long as the cornea, and a long basophthalmite articulating at an angle. Its orbits are narrower than the eyes, but the long podophthalmite and basophthalmite articulate in such a way that allows the eye to be retracted semi-vertically into the orbit. The cornea diameter is on average 5% the carapace length, and is made up of thousands of hexagonal facets packed hexagonally with an approximate diameter of 52 µm (Fig. 9K,L; Table 1).

Finally, extant genera in the subfamily Raninoidinae share the presence of small elliptical eyes on longer eyestalks, all bearing hexagonal facets in hexagonal packing. All three taxa have sub-horizontal orbits with two well-developed orbital fissures and orbital spines. In the studied specimen of *Notopoides latus* (Fig. 10A) the cornea width is 5% the carapace length, and the facets diameter measure around 42 µm (Fig. 10B,C), while in *Notosceles viaderi* (Fig. 10D) the cornea width is less than 3% the carapace length, and the facets measure around 38 µm (Fig. 10E,F; Table 1). The eyestalks of *Raninoides benedicti* (Fig. 10G) are longer than in the other Raninoidinae genera, approximately four times as long as the cornea. The cornea diameter is 2.5% the carapace length, and the facets measure 26 µm in diameter (Fig. 10H,I; Table 1).

#### Cyclodorippoidea

The eyes of cyclodorippoids are little known. The specimen of *Clythrocerus nitidus* studied here (Fig. 10J) has small, roundish eyes, with a cornea nearly as long as it is wide, and as long as the podophthalmite. The cornea diameter is about 11% of carapace length, and is covered in small, well-defined hexagonal facets in hexagonal array. Facet diameter is 35 µm (Fig. 10K, L; Table 1). Previously, round facets in hexagonal packing have been reported for *Krangalangia spinosa* (see Scholtz and McLay, 2009), supporting the absence of mirror eyes in cyclodorippoid crabs.

### Eye preservation in fossil podotreme brachyurans

#### †Callichimaeroidea

†*Callichimaera perplexa* has large globular eyes nearly as wide as long, resting in short eyestalks, and lacks orbits, orbital spines, or any protective structures (Fig. 11). The diameter of the cornea measures approximately 15% of the length of the carapace, and it is covered by small hexagonal to roundish facets in hexagonal packing with an average diameter of 34 µm (Fig. 11C,F; Table 1). One small specimen exhibits a combination of facet shapes and arrays near the junction with the podophthalmite are sub-square, measure approximately 26 µm in diameter, and are packed in a somewhat rectilinear array (Luque et al., 2019a).

#### †Etyoidea

Two specimens of †*Xanthosia* spp. (family †Etyidae) from the Lower Cretaceous (upper Albian, ∼105 Mya) Pawpaw Formation, Washita Group, Texas, USA, show small hemispherical eyes with reduced eyestalks, and a cornea bearing hexagonal facets packed hexagonally (Fig. 12A–F). The diameter of the cornea, preserved in one specimen (Fig. 12C), measures approximately 6% of the width of the carapace, with an average facet diameter of 25 µm (Table 1). The orbits are half as long as the front, and about one-sixth of the carapace maximum width. They are not horizontal but diverge postero-laterally, and have two broad supraorbital fissures separated by a flat, rectangular intraorbital spine (Fig. 12A,B).

One specimens of †*Caloxanthus americanus* (family †Feldmannidae) from the ‘mid’-Cretaceous (Cenomanian, ∼95 Mya) Grayson Formation, Texas, USA, shows round hemispherical eyes resting in reduced eyestalks, and with corneae covered in small hexagonal facets packed hexagonally (Vega et al., 2014) (Fig. 12G–I). The diameter of the cornea measures approximately 10% of the width of the carapace, with an average facet diameter of XX µm (Table 1). As in †*Xanthosia*, the orbits are half as long as the front, and about one-sixth to one-seventh of the carapace maximum width, also diverging slightly postero-laterally.

#### †Necrocarcinoidea

Three specimens of †*Cenomanocarcinus* sp. Stenzel, 1945 (Raninoida: †Necrocarcinoidea: †Cenomanocarcinidae) from the lower-mid Turonian (Upper Cretaceous, ∼90 Mya) and the lower Coniacian (Upper Cretaceous, ∼88 Mya) of Colombia, South America (Fig. 13A–I) exhibit the first and only recorded compound eyes with facets preserved in fossil raninoidans. The Turonian specimens (Fig. 13A–F) bear numerous small hexagonal facets in hexagonal packing. Their eyes rest on small eyestalks, and fit in their short orbits bearing two orbital fissures. The Coniacian specimen, preserved in ventral view (Fig. 13G), preserves its right eye, has a roundish cornea and a short eyestalk (Fig. 13H). As in the Turonian specimens, the eye portion that is exposed bears small hexagonal facets packed in a hexagonal arrangement (Fig. 13I).

#### †Dakoticancroidea

Four specimens of †*Avitelmessus grapsoideus* from the Cenomanian (lower Upper Cretaceous, ∼95 Ma) of Texas, USA (Fig. 14; Table 1). The material examined from the USNM Paleobiology collections preserve very well the cuticle of the overall carapace, but the cuticle of the eye corneae are not preserved. These crab specimens are molts, and that the cuticle of the corneae seems to be eroded away. Many of the eyes in these specimens were still covered by matrix, so the specimens were prepared mechanically to expose fresh eye surfaces in the search of facets. Yet, no facets in †*Avitelmessus* were positively identified (Fig. 14D, H, L), likely because the specimens seem to be exuviae. Although facet shape and packing is still unknown for dakoticancroids, we would expect dakoticancroids to have hexagonal facets in hexagonal packing rather than square facets in orthogonal array. Some dakoticancroids have an orbital bulla reticularis (Bishop, 1984), which was not recognized in the studied †*Avitelmessus* specimens.

### Eye preservation in fossil Eubrachyura

Two fossil eubrachyurans indeterminate from the Santonian (Upper Cretaceous, ∼85 Mya) (Fig. 15A–C) and Campanian (Upper Cretaceous, ∼80–75 Mya) (Fig. 15D–F) of Colombia, have eyes bearing several small hexagonal facets packed in a hexagonal pattern. Likewise, a fossil freshwater crab from the Miocene of Panama (Neogene, ∼16 Mya) (Fig. 15G–I) has three-dimensional eyes bearing minute hexagonal facets in hexagonal packing, but hardly discernible (Fig. 15I).

### Facet shape and packing

Lens packing differs significantly (p < 0.0001) according to facet shape, i.e., all specimens examined that exhibit hexagonal packing also had hexagonal lens shape, and specimens with orthogonal packing possessed square lenses. The morphology of lens packing and facet shape therefore covary in these taxa. As such, principal component analysis shows two distinct clusters representing hexagonal and square lens packing (Fig. 17). PC 1 records the largest amounts of variance (70.5%), representing differences in lens packing. PC 2 (26.8% of the total variance) represents error due to the curvature of the eye, an effect of imaging a curved surface, which affects the distances between landmarks and is uninformative regarding the packing arrangement of ommatidia. The remaining six PCs account for less than 3% of the total variance and do not reveal any major trends in the packing of ommatidia.

## DISCUSSION

The optical mechanisms in ‘intermediate’ podotremes—fossil or extant—are still poorly known, leaving a gap in our understanding of visual systems and ommatidia morphology between the two extremes of the brachyuran tree of life, i.e., the podotreme Homolodromioidea, Dromioidea, and Homoloidea in one hand, and the Eubrachyura in the other hand (Fig. 4). Based on a literature review, plus new data here presented, we examine some aspects of ommatidia packing and facet shape across podotreme brachyurans, and what they can inform about the presence/absence of particular eye types in these groups. In addition, we discuss future directions related to the evolution of the ecology and development of crab eyes, photoreceptors, visual pigments, and vision loss across crabs, highlighting the lack of information available for intermediate crabs.

### Ommatidial packing and facet shape in extant podotreme crabs

External optical features alone cannot reveal details of eye light-path adaptations, however, they are useful for identifying the presence or absence of reflecting superposition eyes by observing their distinctive square facets in an orthogonal lattice (Fig. 2). Eyes of the apposition, refracting superposition, and parabolic superposition types have different internal structural mechanisms to focus the light beams into the retina and form images, but they share the hexagonal to roundish external shape of the facets packed in a hexagonal lattice (Fig. 2). A hexagonal array efficiently packs cylindrical or hexagonal ommatidia into an eye, reducing the angular separation of the ommatidia to a minimum and increasing the eye resolution (Gaten, 1998). Reflecting superposition eyes, however, have distinctive square facets in an orthogonal array indicating ‘mirror’ optics with underlying square crystalline cones.

The eyes of Homolodromioidea, Dromiodea, and Homoloidea all share the plesiomorphic presence of adult eyes with square facets packed orthogonally, typical of reflecting superposition optics. Square facets are essential to the mirror mechanisms (Fincham, 1980; Vogt, 1980, and their presence in homolodromioid, dromiod, and homoloid crabs strongly contrasts with the lack of reflecting superposition eyes in other fossil and extant podotremes and eubrachyurans.

Raninoids or frog crabs are one of the main brachyuran groups for which visual systems are largely unknown, in part due to their cryptic lifestyle and range of bathymetric depths (from 5 to 1000 m depth) (Luque, 2015), making their collection and study difficult. Extant raninoids are adapted for burrowing in sand or soft sediment (Bourne, 1922; Tucker, 1998; Luque, 2015). Their particular ‘frog-like’ morphology with elongated carapace, a pleon that is partially exposed dorsally, elongated mouthparts, modified sternites, naked pleurae, and modified distal podomeres of their walking legs, are regarded as adaptations for their burrowing habit (Luque et al., 2019a). Gaten (1998) suggested that the relatively small eyes in *Ranina* are also an adaptation to a fossorial lifestyle. Extant raninids remain buried in the substratum during the day, emerging at night to search for food (Skinner and Hill, 1986). Some taxa like *Ranilia, Ranina*, and particularly *Cosmonotus*, have relatively large eyes covered in small facets of nearly the same size throughout the cornea, and have long podophthalmite eyestalks that can be held outside the sediment when buried (Fig. 8). Conversely, raninid crabs like *Symethis* (Fig. 7J) show an extreme reduction of the eye and eyestalk, with the cornea enclosed in a reduced orbit and bearing only a couple of hundred facets with different shapes, sizes, and packing (Fig. 7K,L). *Symethis* eyes seem to be degenerate and with poor resolving power.

Since frog crabs occupy an intermediate position between the earliest brachyuran branches (i.e., Homolodromioidea, Dromioidea, Homoloidea), and the more derived groups (i.e., Cyclodorippoidea and Eubrachyura), understanding raninoid optics is essential for testing hypotheses of visual system distributions across brachyurans and their phylogenetic significance. The clear hexagonal facets in hexagonal packing (Figs 8–14), indicate the absence of reflecting superposition in Raninoidea as a whole. The presence of hexagonal facets in hexagonal packing in all raninoid species refutes the assumption by Gaten (1998) (see also table 1 in Porter and Cronin, 2009) that raninoids have mirror eyes just like in basal podotremes (Dromioidea, Homolodromioidea, and Homoloidea). The incorrect inference for raninoids was obviously based on the erroneous supposition that ‘Podotremata’ is a monophyletic group whose members share the presence of reflecting superposition (mirror) eyes. Our findings, on the contrary, indicate that the visual systems present in raninoids are not of the mirror type but either the apposition, parabolic superposition, or refracting superposition type, thus more similar to the visual systems in eubrachyurans and ‘higher’ podotremes (Figs 8–16). Furthermore, Cretaceous stem-group raninoidans such as †*Cenomanocarcinus* (Raninoida: †Necrocarcinoidea) also preserve eyes bearing small hexagonal facets packed in hexagonal arrangement (Fig. 13). †Cenomanocarcinidae belongs to a group of ancient crab-like raninoidans distantly related to †Palaeocorystoidea, which themselves form a grade from which Raninoidea likely evolved (van Bakel et al., 2012; Karasawa et al., 2014; Luque, 2015; Schweitzer et al., 2016; Luque et al., 2019a). This suggests that the loss of mirror optics in adults of the total group Raninoida (i.e., †Necrocarcinoidea, †Palaeocorystoidea, and Raninoidea) most likely occurred in their most recent common ancestor and all of its descendants, more than 90 Mya, which is the age of the fossil cenomanocarcinids with eyes.

The eyes of crabs in the extant superfamily Cyclodorippoidea are also understudied, in part because these crabs are mostly found in deep waters and they have relatively small eyes. In species such as *Krangalangia spinosa*, the eyes are small (perhaps paedomorphic) and the corneal surface is covered by a small number of roundish facets in a hexagonal array (e.g., Scholtz and McLay, 2009, fig. 10). Other cyclodorippoids such as *Clythrocerus nitidus* (A. Milne-Edwards, 1880) have a clear hexagonal pattern of packing and facet shape (Fig. 10J–L). Thus, cyclodorippoid eyes conform with the absence of reflecting superposition eyes with square facets seen in more basal podotremes (except for those species with secondary eye reduction, see below under ‘*vision and eye loss in crabs*’).

### Ommatidial packing and facet shape in fossil podotreme crabs

The morphology of ommatidial packing and facet shape is particularly useful when aiming to understand the fossil record of crustacean compound eyes, since only in a few exceptional cases internal eye structures are preserved (e.g., Vannier et al., 2016). In the enigmatic †*Callichimaera perplexa*, hexagonal to round facets in hexagonal packing are the dominant feature throughout the cornea suggestive of apposition, parabolic superposition, or refracting superposition eyes (Fig.11). One small specimen of †*Callichimaera*, however, has some proximal sub-square facets with rectilinear packing near the contact with the eyestalk (see in Luque et al., 2019a). In decapod crustaceans, the presence of two types of facets in the same eye is uncommon, with only a handful of fossil and extant species showing a combination of hexagonal and square facets. For instance, the larval to early juvenile instars of the shrimp *Oplophorus spinosus* (Brullé, 1839), *Systellaspis debilis* (A Milne-Edwards, 1881), and a post-larval vent shrimp likely of *Rimicaris exoculata* Williams and Rona, 1986, show a mosaic of hexagonal and square facets likely associated with the transition from the larval apposition to the post-larval reflecting superposition eye type (Gaten and Herring, 1995; Gaten et al., 1998). Likewise, a fossil polychelidan lobster specimen from the Jurassic of France (Audo et al., 2019) has two facet shapes in the same eye, but whether they represent a true regionalization of the eye or an artefact of the packing is unclear. The presence of squarish facets in one specimen of †*Callichimaera* thus may be the result of local facet packing rather than a true regionalization or transition between eye types. Furthermore, †*Callichimaera* lacks any associated protective structures such as orbits or orbital spines (Fig. 11), indicating that its eyes must have remained exposed at all times even under times of stress. Such exposed eyes and lack of orbits are mostly seen in crab megalopae, before they metamorphose into their post-larval stage. †*Callichimaera*’s large and unprotected globular eyes and its overall body form have been interpreted as possible paedomorphic retention of larval traits in adulthood (Luque et al., 2019a).

Similar to †Callichimaeroidea, crabs of the superfamily †Etyoidea share with more inclusive podotremes and with eubrachyurans the absence of square facets in orthogonal array, typical of reflecting superposition eyes and present in basal podotreme lineages (Fig. 12). Vega et al. (2014) reported a specimen of †*Caloxanthus americanus* (family †Feldmannidae) also bearing hexagonal facets in hexagonal packing, which together with the specimens of †*Xanthosia* spp. here illustrated confirm the presence of hexagonal facets in species of both etyoid families, unlike the oldest brachyuran groups whose eyes are of the reflecting superposition type. Unlike †Callichimaeroidea and †Etyoidea, for which fossils preserving compound eyes with hexagonal facets have been reported, the corneal eyes of crabs from the superfamily †Torynommoidea are yet to be discovered. Based on our current understanding of the distribution of visual systems across fossil and extant brachyurans, and given the phylogenetic position of torynommoids closer to etyoids and raninoids (Luque et al., 2019a) (Fig. 16), we anticipate the presence of hexagonal facets in hexagonal packing in torynommoid crabs, instead of the square facets in orthogonal packing typical of early brachyuran lineages like dromioids, homolodromioids, and homoloids.

### Eye preservation in other fossil crabs

Although preservation of crab eyes has been considered unusual (Klompmaker et al., 2017), we show that this is untrue. External and internal visual elements in fossil brachyurans do occur in several taxa from different groups, lithologies, and ages, and are underreported in the literature due to biases in recognizing them. Aside from †*Callichimaera*, facet-bearing eyes in fossil crabs have been found in the extinct etyoid †*Caloxanthus americanus* Rathbun, 1935a, from the Cenomanian of Texas (Vega et al., 2014), and in the fossil etyoids, cenomanocarcinids, and the eubrachyuran crabs here reported (Figs 11–14). Tanaka et al. (2009) reported isolated decapod eyes bearing hexagonal facets from the Aptian–Albian Romualdo Formation of Brazil (115–110 Mya), presumably from an achelatan phyllosoma larva, but the systematic placement remains uncertain. Also from the Romualdo Formation are some fossil brachyuran larvae bearing eyes with facets, such as zoeae preserved as stomach contents in the fish †*Tharrhias* (Maisey and Carvalho, 1995; Luque, 2015), and some fossil crab megalopae (Luque et al., 2019b). Both of these fossil crab larvae are modern-looking and have hexagonal facets packed hexagonally, just as modern crab larvae do (Fig. 5). Another fossil crab preserving eyes is †*Ekalakia exophthalmos* Feldmann et al., 2008 (Dromiacea: †Glaessneropsidae), from the Campanian–Maastrichtian (70 Mya) Pierre Shale Formation in the USA. Unfortunately, no corneae or facets are preserved in any known specimen. Miocene grapsoid crabs preserved in amber from Mexico (Serrano-Sánchez et al., 2016) also preserve facets, most likely hexagonal as in all other grapsoids. Overall, the fossil record indeed has several remarkably preserved eyes wherein ommatidia arrangements are available for interpretation. Considering the utility of this character for resolving phylogenies, and the ability to discriminate the character using unbiased morphometrics (Fig 17), further effort is warranted in cataloguing fossil ommatidia shapes.

### Phylogenetic implications

Although decapod larvae have apposition eyes (the simplest type of compound eye), adult decapods of most shrimp, lobster, galatheoid anomuran, and early podotreme brachyuran clades share a unique reflecting superposition visual system not seen in other crustaceans outside Decapoda (Land, 1976; Scholtz and McLay, 2009; Tudge et al., 2012; Gaten et al., 2013). Noticeably, many decapods have independently retained larval apposition eyes (Gaten, 1998; Cronin and Porter, 2008; Porter and Cronin, 2009); or have evolved refractive or parabolic superposition eyes while retaining hexagonal facet shape and packing (Nilsson, 1988). Apposition and reflecting superposition eyes are more or less homogeneous across taxa, while the optical mechanisms in the parabolic and refracting superposition are more variable and with intermediate forms (Nilsson, 1983; Porter and Cronin, 2009).

Reflecting superposition eyes likely evolved only once in a recent common ancestor of Decapoda during the Paleozoic. Among crabs, mirror optics are found in the earliest brachyuran lineages i.e., Homolodromioidea, Dromioidea, and Homoloidea, but are absent in Raninoidea, Cyclodorippoidea, and the most anatomically diverse and species-rich group of crabs: the Eubrachyura (Fig. 14). We should keep in mind, however, that ecology plays a crucial role in shaping the visual systems in an organism to better suit their biology, and a number of taxa with a given eye type—or lack thereof—are likely to undergo adaptations to better suit their current ecological pressures (see below).

### Other aspects of crab vision and future directions

#### Ecology and development of crab visual systems

For any given species, living or fossil, the anatomy of the eye partly reflects selection pressures from the required visual tasks that must be performed within the light environments in which it lives. As in most crustaceans, brachyuran life history progresses through several stages living in different habitats, and therefore many aspects of the visual system (facet shape, optics, visual pigments) may change throughout ontogeny. The early larval stages are generally pelagic and have transparent apposition eyes (Fig. 5). Larval apposition optics are specialized for open water habitats, and are thought to provide as much camouflage (predator avoidance) to the larva as possible by decreasing the pigmented portion of the retina to minimize its diameter. This results in a ‘clear zone’ between the cones and the rhabdom, thus the larval apposition eye functionally mimics a superposition eye (Gaten, 1998; Feller and Cronin, 2014). At metamorphosis into the juvenile stage, these larval eyes are remodeled to form the diversity of compound eye types found in adult crabs, including reflective superposition optics in podotreme brachyurans (Gaten, 1998), and parabolic superposition optics in families of Eubrachyura (Nilsson, 1988). Conversely, crabs that inhabit consistently similar habitats throughout their life cycle do not seem to change their visual sensitivity during metamorphosis (Cronin et al., 1995). For reflecting superposition eyes (Fig. 2C, 5), note that the ommatidial facets are square as this is the only shape that can reflect light onto the retina.

Variations in the arrangement of eyes among species reflect specializations related to visual tasks within a given environment. Crabs that live in relatively flat environments (e.g. Dotillidae, Goneplacidae, Heloeciidae, Ocypodidae, Mictyridae, Macrophthalmidae) tend to carry their eyes close together at the end of long eye stalks, or occasionally midway along an elongated eye stalk (as in the horned ghost crab *Ocypode ceratophthalma*). These taxa may have a need to resolve the distance of objects in their flat environments (Zeil et al., 1986). Crabs living on rocky shores or in mangrove forests (e.g., Grapsidae, Sesarmidae) tend to carry their eyes far apart on short eye stalks (Zeil et al., 1986; Zeil et al., 1989; Zeil and Hemmi, 2006; Davie et al., 2015). Most occurrences of elongated eyestalks in brachyurans are from the Eubrachyura, although examples can also be found in groups such as the Latreillidae and Homolidae (Fig. 7), likely the result of different selective pressures to those associated with flat visual environments (Davie et al., 2015). There is no close phylogenetic relationship among groups with elongated eyestalks.

#### Vision and eye loss in crabs

Although most crabs have functional eyes throughout some or all of their growth stages, from light-sensing to image forming, dramatic secondary reductions or complete losses of eyes has happened many times independently in crabs, and cannot be expected to inform phylogenies at a high taxonomic level. These instances are generally limited to small groups of species that live in dark habitats such as the deep sea and caves. Species living in caves exhibit a range of eye reductions, and include representatives from Hymenosomatidae (e.g., Ng, 1991; Husana et al., 2011), Grapsidae (e.g., Ng et al., 1994), Potamidae (e.g., Yeo and Ng, 1999; Ng, 2017; Gecarcinucidae (e.g., Stasolla et al., 2015), and Parathelphusidae (e.g., Takeda and Ng, 2001; Husana et al., 2009), among others. In smaller individuals of *Cancrocaeca xenomorpha* (Hymenosomatidae), the eye seems completely absent (Ng, 1991; Ng and Chuang, 1996), although larger individuals have discernible remnants of the eyes.

In the deep sea, many crustaceans have responded to low levels of light by either increasing eye size and visual sensitivity, or by undergoing reduction in visual structures leading to vestigial eyes. As an example of eye reduction, the deep-sea crab *Cymonomus bathamae* Dell, 1971 (Cyclodorippoidea: Cymonomidae) has eyes that externally lack corneal facets, crystalline cones, and pigment cells, and the rhabdoms are irregular in shape and cell number (Chapman, 1977). Deep-sea hydrothermal vent crabs of the family Bythrograeidae like *Austinograea williamsi* Hessler and Martin, 1989 and *Bythograea thermydron* Williams, 1980 (see also Guinot, 1990) represent an extreme case of eye loss in crabs. In this group, the larval stages have limited pelagic capability, and the adults are completely benthic on the substrate adjacent to the vents. Larvae have conventional apposition compound eyes, which regress in the adults to form a featureless mass of tissue lacking cornea or lenses, and specialized for detecting the dim, red light emitted from hydrothermal vent water (Jinks et al., 2002). The adult vestigial tissue is located at the base of the second antenna, while the orbits have become reduced and shallow as they serve no protective purpose for the eye anymore.

#### Photoreceptor and retinal structure

Photoreceptor cells are responsible for transduction of visual signals, and in arthropod compound eyes are clustered together as ommatidia. Thus far, the photoreceptor arrangement has only been studied in eubrachyurans and other non-brachyuran decapods which possess mirror eyes, but not in podotremes. Eubrachyuran compound eyes in general follow the crustacean plan, containing eight photoreceptor cells (retinular cells) that fall in two anatomically distinct classes. Seven of the retinular cells (R1-7) contribute microvilli to a main receptor, that itself sits below a receptor formed from the final, single retinular cell (R8, located distally within the rhabdom). These two receptor types are generally tuned to absorb medium and short wavelengths from the blue-green (R1-7) and violet or ultraviolet (UV) (R8) portions of the spectrum. Based on this photoreceptor anatomy and physiological measurements of receptor sensitivity (Cronin and Forward, 1988; Jordão et al., 2007), most crabs have been hypothesized to be functional dichromats (Horch et al., 2002). However, both anatomical and molecular studies suggest as of yet uncharacterized potential diversity in brachyuran receptor morphology and sensitivity. In one of the only such studies conducted, expression of visual pigments in the Atlantic sand fiddler crab, *Uca pugilator*, found cell specific expression, suggesting functional diversity at the level of individual retinular cells (Rajkumar et al., 2010; further discussion below).

Anatomically, studies of the microvilli arrangement in fiddler crab receptors R1-7 have also suggested fine tuning (maximal separation of polarization contrast sensitivity) both regionally across the retina, and longitudinally across the length of each photoreceptor (Alkaladi et al., 2013). These anatomical studies suggest the potential for fine scale diversity within photoreceptors R1-7 that will affect overall chromatic and polarization sensitivity among species, but requires studies across many more species to fully characterize patterns relative to phylogeny and ecology.

#### Visual pigments and phototransduction

From a molecular perspective, genetic and genomic methods have long advanced the field of vision research. However, our current knowledge of the genes involved in vision and phototransduction, i.e., the conversion of light to electrochemical signals, is severely limited across decapod crustaceans (Henze and Oakley, 2015; Ramos et al., 2019; Zhang et al., 2019), with all crab data restricted to only a few eubrachyurans. Nevertheless, visual pigments play a critical role in the activation of the phototransduction signaling cascade, and consist of an opsin protein bound to a chromophore (Pérez-Moreno et al., 2018). To date, the few studies that use molecular approaches aim to characterize the visual pigments, and more specifically opsin genes, present in the retina of the eyes and/or eyestalk. Yet those few impactful studies from the past three decades have advanced our knowledge of vision across eubrachyurans and lay the groundwork for future integrative research that combines genomics with other approaches, and that will investigate podotreme genomic data.

One of the first studies of vision in brachyurans, and across crustaceans was carried out by Sakamoto et al. (1996), who aimed to test the hypothesis that crustaceans had multiple visual pigments in their retinas. Mounting evidence from spectral sensitivity studies suggested that decapods and stomatopods had multiple color receptors, and behavioral studies indicated fiddler crabs could discriminate colors (Hyatt, 1974, 1975). To test for multiple visual pigments, Sakamoto et al. (1996) used cDNA sequencing in the shore crab, *Hemigrapsus sanguineus,* to successfully isolate two different opsin sequences, BcRh1 and BcRh2, and to investigate opsin expression via *in situ* hybridization. Two opsins were expressed in similar levels across all seven retinula cells (R1-7) that form the main rhabdom in each ommatidium. No definitive signal of opsin expression was observed via *in situ* hybridization in the small distal retinular cell (R8) (Sakamoto et al., 1996), possibly due to the cell’s smaller size (Stowe, 1980), or lack of an appropriate probe for a UV opsin. Visual sensitivity was further investigated using electroretinogram (ERG) measurements, suggesting the eye of *H. sanguineus* has a maximum sensitivity of 480nm. In the ERG, a slight shoulder was detected at 330-400nm possibly indicating a second shorter-wavelength sensitivity, however this could not be confirmed with genetic results. It is plausible this signal was associated with the R8 cell, and possibly contained a third putative opsin sensitive to UV-wavelengths. Overall, this was one of the first molecular studies to demonstrate that two opsin proteins were present in one photoreceptor of the retina of the crab, *H. sanguineus,* with maximum absorbance in the blue-green range.

Over a decade later, Rajkumar et al. (2010) used molecular methods to investigate the spectral properties of fiddler crab eyes after behavioral (e.g., Hyatt, 1974, 1975; Detto, 2007; Detto and Backwell, 2009) and physiological studies (e.g., Hyatt, 1974; Scott and Mote, 1974; Horch et al., 2002; Jordão et al., 2007) provided contradictory evidence for the number of visual pigments in the retinular cells. Using cDNA sequencing methods, Rajkumar et al. (2010) successfully characterized three distinct opsins (UpRh1, UpRh2, and UpRh3), from *Uca* (*Leptuca*) *pugilator*. *In situ* hybridization was used to examine expression levels in all eight retinula cells, and phylogenetic annotation was used to assign putative spectral sensitivities. UpRh1 and UpRh2 were expressed in the R1-7 cells, nested within a clade of arthropod opsins sensitive to middle wavelengths (MWS). Interestingly, UpRh1 was only expressed in five cells whereas UpRh2 was expressed in three, indicating co-expression of these opsins occurs in one retinular cell. UpRh3 was expressed only in the R8 photoreceptor cell, and was phylogenetically placed within a clade of arthropod opsins sensitive to short wavelengths (UV). Therefore, *U. pugilator* has the opsin repertoire to discriminate color with possible trichromatic vision, if UpRh1 and UpRh2 have different peak absorbances. At the time of writing, the opsin sequences and spectral sensitivities are only known from two species of brachyuran crabs, *U. pugilator* and *H. sanguineus.* Subsequent sequences from a handful of other species (restricted to eubrachyurans) are now available on NCBI, although none are linked to expression studies targeting the eyes in crabs.

To date, genomic investigations of brachyuran visual systems are lacking. While a handful of studies have used RNA-seq to characterize the genes expressed within eyes or eyestalks, understanding the visual capabilities of crabs was not the main objective of this research (Hui et al., 2017; Lv et al., 2017; Yingdong et al., 2019). For example, tissue-specific RNA-seq from the eyestalks of *Portunus trituberculatus* and *Eriocheir sinensis* identified genes involved in molting and the circadian cycle, respectively. The data from *E. sinensis* are especially useful due to the publication of a draft genome sequence (Song et al., 2016). A separate study extracted RNA from the vestigial eyes of the hydrothermal vent crab, *Austinograea alayseae* to investigate genes associated with living in extreme environments (Hui et al., 2017). From the vent crab, evidence was found for all genes involved in the fly phototransduction pathway and two distinct opsin sequences, however the expression levels were low. Although these studies did not focus on visual ability, these data could be used in combination with more targeted approaches to advance the field of brachyuran vision.

Significantly, the presence of opsin sequences in the genome alone does not reveal anything about functionality, however these data lay the foundation for inferring the visual ability of any animal taxon. Integrative approaches that combine genomic, gene expression, physiological, microscopy, behavioral, and eventually gene editing studies, have the potential to revolutionize our understanding of vision across metazoans, however these approaches are still in their infancy across brachyurans, and data are entirely lacking for podotremes (the group with the most interesting and potentially phylogenetically informative variation in ommatidial morphology). As genomic data are generated for additional crab species, visual pigments should be targeted by eye-specific transcriptomics and/or the design of genomic probes based on previous visual pigment annotations (Speiser et al., 2014; Schott et al., 2017; Pérez-Moreno et al., 2018). Tracing the evolution of photoreceptor genes and eye morphology across a robust crab tree could be an informative first step to reconstructing the evolutionary history of vision across this iconic group.

## CONCLUSIONS

Early brachyuran clades like the podotremes Dromioidea, Homolodromioidea, and Homoloida have reflecting superposition ‘mirror’ eyes, characterized by square facets packed in a rectilinear lattice. Mirror eyes are plesiomorphic for Decapoda, and are found in most shrimps, lobsters, several anomurans, and the least inclusive brachyuran clades mentioned above. This supports the view that mirror eyes were present in the most recent common ancestor of crown group brachyurans.

Conversely, closer ingroup podotreme lineages like †Callichimaeroidea, †Etyoidea (i.e., †Feldmannidae, †Etyiidae), †Necrocarcinoidea (e.g., †Cenomanocarcinidae), Raninoidea and Cyclodorippoidea, together with the so-called ‘higher’ true crabs or Eubrachyura, lack mirror eyes, which may have been lost in a most recent common ancestor for those groups. The expression of eyes with hexagonal/roundish facets in hexagonal packing see across adult “higher” podotremes and eubrachyurans can be interpreted as a result of the secondary, paedomorphic retention of larval apposition eyes, or exaptation of a pre-adapted larval apposition eye to function as parabolic superposition eyes (Porter and Cronin, 2009). We conclude that the retention of apposition eyes in ‘higher’ podotremes and eubrachyurans has existed since at least the Early Cretaceous, more than 100 million years ago (Fig. 16). The distribution of eye types among brachyuran crabs provides evidence to refute the monophyly of podotremes, instead suggesting the groups that share apposition eyes form a paraphyletic grade with eubrachyurans, consistent with recent molecular and morphological phylogenetic works.

Ecology appears to be an important driver of visual systems among higher taxa, especially with respect to terrestrialization, colonization of fresh water, diurnal activity, specific dietary habits, and bathymetry, potentially leading to the independent loss of mirror eyes in many taxa. Ongoing work aims to shed light on whether predictable “rules” account for the potential convergent origins and/or losses of apposition and mirror eye types among true crabs through time.

## Acknowledgements

J.L. thanks Rafael Lemaitre, Karen Reed, Mark Florence and Conrad Labandeira (USNM, USA) for providing access and loaning specimens from the Invertebrate Zoology and Paleobiology collections at the Smithsonian Institute. Peter Davie (QWM, Australia), for help accessing the QMW Invertebrate Zoology collections at the Queensland Museum, Brisbane, Australia. To Daniéle Guinot, Paula Martin-Lefebre, and Laure Corbaire (MNHN Paris, France), for help accessing the Palaeontology and Invertebrate Zoology collections at the Muséum national d’Histoire naturelle, Paris, France. To Eric Lazo-Wasem and Lourdes Rojas (Yale University, USA) for export permits and photography of specimens from the Yale Peabody Museum Invertebrate Zoology Collections. To Jody Martin (LACM, USA) for facilitating the examination of extant crab eye samples of specimens deposited at the YPM and the LACM collections. To Michelle Pinsdorf (Department of Paleobiology, USNM) for help with the preparation of the USNM specimens of †*Avitelmessus grapsoideus*, and to Liath Appleton and the Non-vertebrate Paleontology Lab at the Jackson School Museum of Earth History (University of Texas) for images of †*Caloxanthus americanus.* John Douglass provided comments on early drafts of the manuscript. Jorge Ceballos (STRI, Panama), Arlene Oatway and Nathan Gerein (UofA, Canada), and Elissa Martin (Yale Peabody Museum) provided SEM assistance. Marcela Gómez, Mauricio Pardo, and the Colombian Geological Survey supplied the export permits of the Colombian fossils here studied.

## Funding

Partial funding for this study was provided to J.L. by the Smithsonian Tropical Research Institute Short–Term Fellowship Program (STRI-STF) (Panama), the American Museum of Natural History (AMNH) Lerner Gray grant (USA), the Sedimentary Geology, Time, Environment, Paleontology, Paleoclimatology & Energy (STEPPE) Student Travel Award Grant (USA), the Fondo Corrigan-ACGGP-ARES (Colombia), and the Natural Science and Engineering Research Council of Canada Graduate Scholarship (NSERC CGS–D). Additional support was provided by the Izaak Walton Killam Memorial Scholarship, the Andrew Stewart Memorial Graduate Prize, the University of Alberta President’s Doctoral Prize of Distinction, the Devendra Jindal Graduate Scholarship, the Alberta Society of Professional Biologists Graduate Scholarship, and the Kay Ball Memorial Graduate Student Research Travel Award (Canada), and the Natural Science and Engineering Research Council of Canada Postdoctoral Fellowship (NSERC PDF). Other support to J.L. via the Yale Institute for Biological Studies (YIBS), and NSERC Discovery Grants RGPIN 04863 to A.R.P. (Canada) and RGPIN-2014-06311 to W.T.A. (Canada). This work was also supported by the National Science Foundation DEB #1856679 to J.M.W. (USA).

## Competing interests

The authors declare that they have no competing interests.

## Data and materials availability/ accessibility

All data needed to evaluate the conclusions in the paper are present in the paper.

